# Comparison of multiple video tracking-based behavioral summary approaches for compound discrimination

**DOI:** 10.64898/2026.07.20.739643

**Authors:** Marti Ritter, Serena Deiana, Alina Ritter, Carsten T. Wotjak, Michael Brecht, Amarender R. Bogadhi

## Abstract

The rapidly increasing number of video tracking-based behavioral summary tools and methods raises the question as to the most suitable approaches for pharmacological fingerprinting in pre-clinical research. We have recently shown that social context has a strong effect on behavioral syntax in mice, further suggesting that treatment effects can be context dependent.

Here, we aim to answer the question whether there is an optimal combination of context and behavioral summary method for effect detection and discrimination of different psychoactive substances in a controlled environment. To this end, we applied eight different treatment-dosage pairs (amphetamine 1.5,3,6 mg/kg; modafinil 5,10,50 mg/kg; seltorexant 3,10 mg/kg) and evaluated five different approaches to behavioral summary: parametric aggregation, unsupervised segmentation in Keypoint-MoSeq (KPMS) & Variational Animal Motion Encoding (VAME), and supervised segmentation in Simple Behavioral Analysis (SimBA) & A-SOiD, across two different contexts (Solitary & Social) in 314 recordings of freely moving mice in an open-field arena.

Surprisingly, our results show no significant differences in performance across models and context. Across treatment effect detection to treatment-dose discrimination, all models showed performance significantly above chance that was insensitive to various data limitations and extensions. Overall, our study shows that under the tested conditions and treatments, the choice of a behavioral summary model does not meaningfully affect the description of treatment effects. Simple aggregate measures from tracking data and machine learning based behavioral summary approaches that are expensive, in terms of training data and computational resources, performed equally well.

Our findings taken together with literature suggest that the fuller decomposition of complex behavior through unsupervised machine learning might be necessary for the description of large-scale datasets but does not necessarily align with the goals present in smaller-scale treatment discrimination tasks common in pre-clinical research.

## 1 Introduction

Pre-clinical research for drug discovery relies heavily on studies of disease-related circuit-to-symptom associations, often in mice. Animal experiments can reveal the foundational aspects of complex behaviors but require accurate measurement and description of observable behavior and intervention effects. Traditionally, mice were used in comparative psychology in specialized but reductive tasks, such as three-chamber social approach tasks (Nadler et al. 2004), or tube tests to determine hierarchies (Fan et al. 2019), in order to evaluate complex social behaviors with fixed, reproducible metrics (Bordes et al. 2023). This enabled an easy, reliable measurement of treatment effects, but also limited the type of effects that could be observed (Peleh et al. 2019). To allow a more natural, complete range of behaviors one could investigate mice in an unconstrained and more naturalistic environment, but this still required the parametrization of complex behavior into quantifiable metrics, such as counting entries into tunnels (Shillito 1970) or beam breaks (Beninger et al. 1985).

To avoid the risk of leaving many aspects of naturalistic behavior unrecognized, it was necessary to develop more advanced tools (Von Ziegler et al. 2021). By applying computer vision approaches, such as the segmentation of mouse outlines from video recordings, it became possible to track locomotion parameters automatically in an open field arena (Prut and Belzung 2003). Many integrated systems became available that allowed the automated extraction of body center position and movement direction (Lim et al. 2023) to detect genetic and pharmacological effects (Fraser et al. 2010). While this description is reproducible, detailed, and high-throughput, it also loses the connection to established ethological labels for characteristic mouse behaviors (e.g. grooming, rearing, and exploration), so called “ethograms”, which were part of the earliest descriptions (Abeelen 1964).

Marker-less pose tracking software allowed the automated extraction of a large amount of pose information available in the already existing ethological video datasets. DeepLabCut (Mathis et al. 2018) or SLEAP (Pereira et al. 2022), for example, also enabled the capture of nuanced treatment effects, and were a valuable foundation for advanced machine learning approaches. Many of the most common natural behaviors (grooming, rearing, locomotion) have been successfully modeled in supervised machine learning pipelines (Goodwin et al. 2024). While this approach holds the potential to differentiate between therapeutic drug classes and mouse strains (Brodkin et al. 2014), it is suffering from limited reproducibility due to a lack of standardized definitions for common behaviors, and requires labeled behavioral, sometimes context specific, training data on top of the extracted tracking data. To mitigate the latter issue, active learning pipelines have been developed (Tillmann et al. 2024).

While these supervised approaches are a move away from previous reductionist assays, the limitation to specific, subjectively pre-defined behavioral patterns is a disadvantage of supervised descriptions of complex behavior (Gris et al. 2017). In recent years novel approaches to unsupervised behavior decomposition have arisen. These tools use short recurring sequences as descriptors of complex behaviors, enabling researchers to find previously hidden behavioral fingerprints of disorders, such as epilepsy (Gschwind et al. 2023), and explore the effects of treatments in a very high-dimensional behavioral space (Wiltschko et al. 2020). A variety of these tools exist at this point, able to use three dimensional recordings (Lin et al. 2024) and body-point representations (Hsu and Yttri 2021; Luxem et al. 2022). Most of them are fully unsupervised, with user input limited to hyperparameters, and some allow input to control motif timescales (Weinreb et al. 2024).

Across all basic behavioral functions required for survival, social interactions are some of the most complex and have been a focus of ethological investigations, with many specialized systems built to track related behaviors (Peleh et al. 2019). Recent research done in our lab (Ritter et al. 2025) revealed that unsupervised behavior segmentation using Keypoint-MoSeq produces syllables whose expression is strongly affected by the presence of a conspecific, even though these motifs do not directly relate to previously established ethological labels for “social” behaviors. We hypothesize that social context could improve the dynamic range for better discrimination of treatment effects, as social behaviors are highly specialized and sensitive to disturbances.

In this study, we aimed to test this hypothesis across a range of behavioral summary models: Parametric aggregation, supervised behavior detection with SimBA (Goodwin et al. 2024) and A-SOiD (Tillmann et al. 2024), unsupervised motif detection with Keypoint-MoSeq (Weinreb et al. 2024) and VAME (Luxem et al. 2022), to determine the optimal combination of behavioral description and context for differentiation within a set of treatments and dosages.

In the simplest case of parametric aggregation, we computed parameters such as speed and heading (see Methods), and in the case of machine learning-based methods it is possible to detect complex behaviors based on latent features found in raw data either without manual annotations (unsupervised machine learning, KPMS & VAME) or with manual annotations (supervised machine learning, SimBA & A-SOiD), which help guide the output to previously established definitions of behavior (see “ethograms”).

As number and nature of the parameters returned by the behavioral summary methods differ widely, we chose to evaluate performance with approaches insensitive to the dimensionality of the output. All tests were performed on three analytical levels corresponding to tasks encountered in drug differentiation studies: The comparison to a vehicle to detect an effect, the comparison between drugs to identify the treatment, and finally the comparison of drug-dosage pairs to fully discriminate treatments. We administered amphetamine, modafinil, and the orexin 2 receptor antagonist seltorexant in different doses and normalized their effects to a common control group. We expected the two stimulants (amphetamine and modafinil) to exert opposite effect to Seltorexant (Bonaventure et al. 2015), given that orexin 2 receptor antagonists are developed for the treatment of hyposomnia (Muehlan et al. 2020). The separation of amphetamine (Yates et al. 2007) and modafinil (Simon et al. 1996), in contrast, has been shown to be potentially difficult in prior literature (Wiltschko et al. 2020), since both show stimulating effects, but with different mechanisms of action.

Surprisingly, we found no significant differences in performance across models and contexts for the treatments we chose. All evaluations performed significantly above chance, were able to detect social context, and were insensitive to a variety of post-hoc data limitations and extensions. This suggests that, within the conditions present in this study, higher-dimensional behavioral features from machine learning-based methods did not significantly add to the description of treatment effects when compared to the simple parametric aggregation method. Additionally, behavioral changes in social context did not significantly enhance the detection of treatments in our study.

## 2 Materials and Methods

### 2.1 Animals

157 male C57BL/6J mice (∼5 weeks old at time of experiment) were obtained from Janvier Labs. They were housed in groups of 4 and kept in a light- and humidity-controlled facility (regular light cycle). Along with the treated animals we also used the same number of mice from a different cohort (yet same age, sex, and source) as stimulus animals (see Experimental procedure). The stimulus animals were kept novel to the treated animals and housed in an isolated facility. All procedures followed the regulations for animal experimentation enforced by the local district administration’s animal welfare commissioner of the state of Baden-Württemberg.

### 2.2 Open Field Arena Setup

The recordings were performed in a square arena (453mm Width x 453mm Depth x 400mm Height) with walls made of clear plexiglass and a removable gray plastic floor to facilitate cleaning (see Suppl. Figure 1). It was equipped with a dimmable LED strip and an analog camera (768px x 576px, 25.23Hz, grayscale, Weldex WDH-2500BS). A wooden enclosure surrounded the arena on all sides, except for the front, which was covered by a red plexiglass door. Throughout the recording the arena was illuminated at a stable level of around 150lux. Two smaller, circular enclosures (Mouse Grid Cage by Ugo Basile, Product-Nr. 46523-003) made from gray plastic with a diameter of 90mm and a height of 215mm were placed in the two upper corners of the arena.

### 2.3 Treatment

All animals were injected either orally (PO), subcutaneously (SC), or intraperitoneally (IP) with one compound and dosage each. Used compounds were seltorexant (PO; 3 & 10 mg/kg), amphetamine (SC; 1.5, 3, 6 mg/kg), and modafinil (IP; 5, 10, 50 mg/kg). Group sizes were different between treatment groups and vehicle groups (see Table 1).

**Table 1.** Compounds, dosages, administration routes, and group sizes.

| Compound | Dosage [mg/kg] | Assigned Dosage Term | Administration Route | # of Mice |
| --- | --- | --- | --- | --- |
| Amphetamine | 1.5 | Low | Subcutaneous | 16 |
|  | 3.0 | Medium | Subcutaneous | 16 |
|  | 6.0 | High | Subcutaneous | 15 |
| Modafinil | 5.0 | Low | Intraperitoneally | 15 |
|  | 10.0 | Medium | Intraperitoneally | 16 |
|  | 50.0 | High | Intraperitoneally | 16 |
| Seltorexant | 3.0 | Low | Orally | 15 |
|  | 10.0 | High | Orally | 16 |
| Vehicle |  | Medium | Intraperitoneally | 8 |
|  |  | Medium | Orally | 8 |
|  |  | Medium | Subcutaneous | 16 |

These treatments were chosen to allow testing for the discriminability of opposing effects, such as between stimulants (amphetamine and modafinil) and orexin receptor antagonists (seltorexant, used in the treatment of i.e. insomnia) (Muehlan et al. 2020; Bonaventure et al. 2015), and stimulating effects with differing mechanisms of action, as found between amphetamine (Yates et al. 2007) and modafinil (Simon et al. 1996). In addition, prior literature showed that differentiation between modafinil and amphetamine is potentially difficult (Wiltschko et al. 2020).

The specific vehicle used in the control group depended on the treatment group, with a matching administration route: amphetamine (SC) was matched to a control group treated with NaCl (SC), modafinil (IP) was matched with 0.5% Na-CMC/Tween-80/NaCl [ml] (IP), and seltorexant (PO) was matched with 20%HPßCD in McIlvaine (PO).

All injections were performed 20 minutes before the recordings started.

### 2.4 Experimental procedure

To reduce stress during the experiment all stimulus animals were habituated to cubicles by placing them inside a cubicle twice for 15 to 20 minutes each in the week before the experiment. The experiments always began between 7:30 and 10:45 am. An overview of the recording and procedure timeline can be found in Figure 1A.

**Figure 1.**
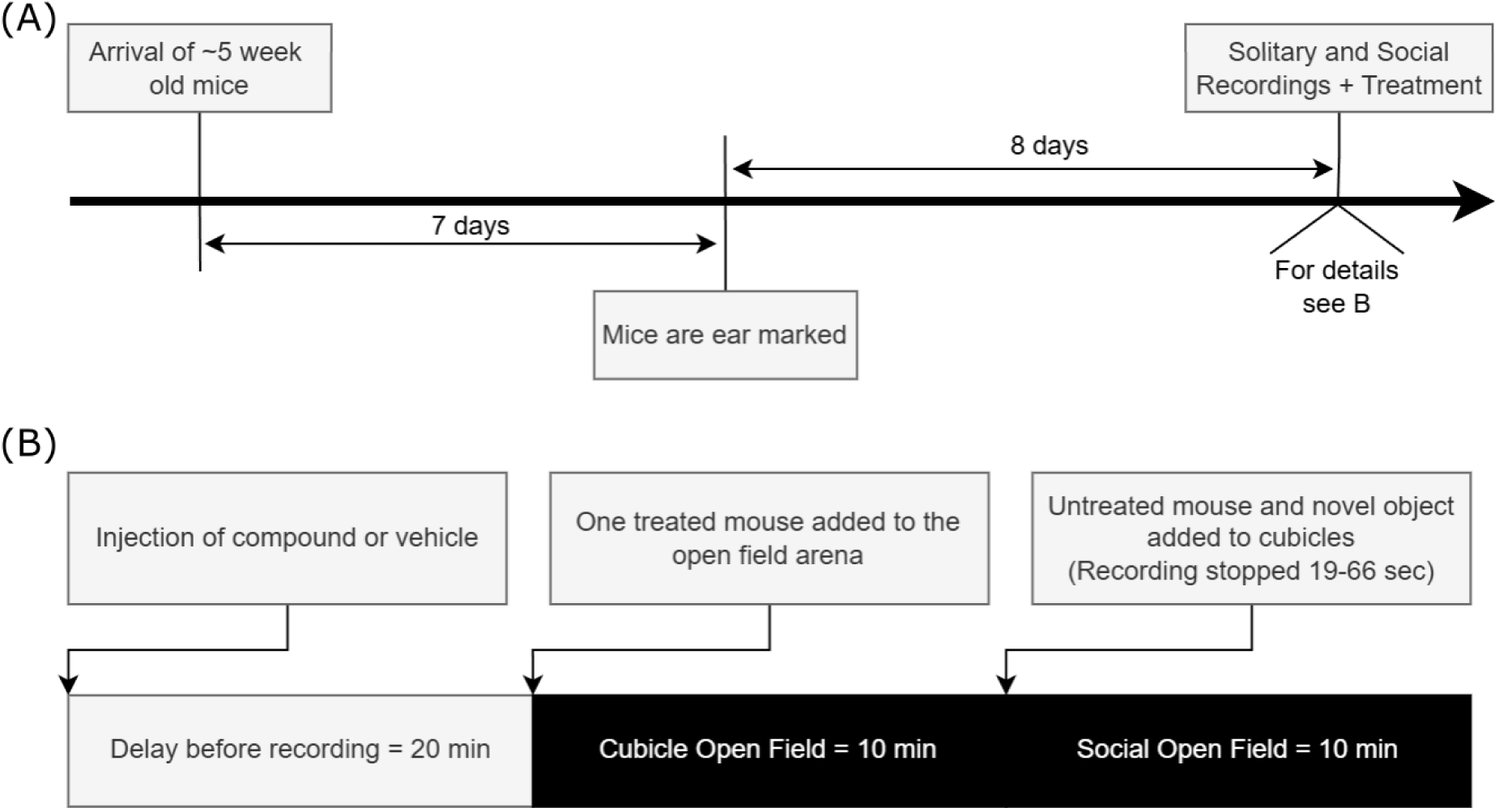
Experiment, procedures, and recordings timeline. (A) ∼5-week-old mice were ordered from Janvier and ear marked a week after delivery. Roughly one week later the recordings were performed. (B) Treatment was applied to mice through varying routes, and after 20 minutes had passed a single mouse was placed in the open-field arena, which already contained two empty containers. This initial recording lasted for ten minutes before a short pause, during which another previously unknown, untreated mouse and an object were placed in the containers. Afterwards, another 10-minute recording was performed.

After application of the treatment or vehicle, the mice returned to their home cages for 20 minutes. Afterwards the mice were transferred to the recording setups and were video recorded for another 20 minutes. A short break (19-66 seconds, avg. 32 seconds) occurred in the middle of the recording, as another (untreated) mouse and a novel object were added to the cubicles contained in the arena. All recordings were controlled manually through Stoelting Any-Maze software (version 7.4). For an example timeline for a single recording session see Figure 1B.

### 2.5 Data collection

#### 2.5.1 Stoelting Any-Maze

We used Stoelting Any-Maze (Figueiredo Cerqueira et al. 2023) version 7.4 to record metadata and videos (no additional tracking parameter extraction) in our study. Further analysis of the resulting videos revealed short breaks of various lengths (1 to 47 frames per occurrence, 1 to 773 cumulative frames in 1 to 42 separate occurrences per video) in a small subset of videos (46 out of 320, ∼14.4%). As each video was 10 minutes long, this affected only a small part of the total dataset (10921 frames / 7.21 minutes out of roughly 4844160 frames / 53.3 hours, ∼0.23%). Relatively more frames were affected in the Vehicle and seltorexant datasets (∼0.34% and 0.42% respectively) compared to amphetamine and modafinil recordings (∼0.09% and ∼0.16% respectively). We resolved this issue by removing affected videos entirely from the training dataset and only applying trained models to the full dataset (including affected videos).

#### 2.5.2 Training dataset

We trained all supervised and unsupervised models only on recordings of vehicle-treated mice. Due to the above-mentioned issue of dropped frames, we decided to limit the training set from a potential 64 videos (32 vehicle mice with 2 phases each, 10.7 hours) to 49 unaffected videos (∼8.2 hours). This decision was made to ensure that the unsupervised models were not affected by the occurrence of unexpected static frames in the dataset. All videos were used during the inference and downstream analysis.

#### 2.5.3 SLEAP

We used SLEAP (Pereira et al. 2022) to track a set of 12 key points on the body of the freely moving mouse. This labeling schema was adapted from Weinreb et al. (2024). For a more detailed description of the SLEAP model used in this approach, see Ritter et al. (2025). As the various behavioral summary models used in this study required different input formats, we translated this dataset using the tools included in the SLEAP Python library. In the case of the two supervised methods, this also required the truncation of four key points, to match this schema to the default used in SimBA, which was also applied to A-SOiD (see Supplementary Methods, Suppl. Figure 2A).

#### 2.5.4 BORIS

We used BORIS (Friard and Gamba 2016) Version 9.0.6 to manually curate the base dataset for supervised learning methods. For this purpose, we sampled 36 out of 49 available training videos across phases, sessions, and subjects.

All videos were manually annotated with a set of 6 behaviors: Climbing, grooming, rearing, and three types of sniffing (at the cubicle, towards the content of the cubicle, and in the environment). We utilized the exclusion matrix functionality included in BORIS and excluded logically contradictory events from co-occurring, in addition to the manual curation (see Supplementary Methods).

### 2.6 Behavioral summary models

As an overview over the possible approaches to behavioral summary, we selected a total of 5 approaches: one purely parametric aggregation into scalar values (e.g. distance moved or relative distance, see Suppl. Figure 2B and C), and two unsupervised as well as two supervised behavior segmentation models, which detected the occurrence of bouts, recurring patterns of behavior of variable length. For simplicity’s sake we will refer to all of these approaches as “behavioral summary” (Wiltschko et al. 2020) models in this manuscript. For an overview of all possible processing flows, see Figure 2A.

**Figure 2.**
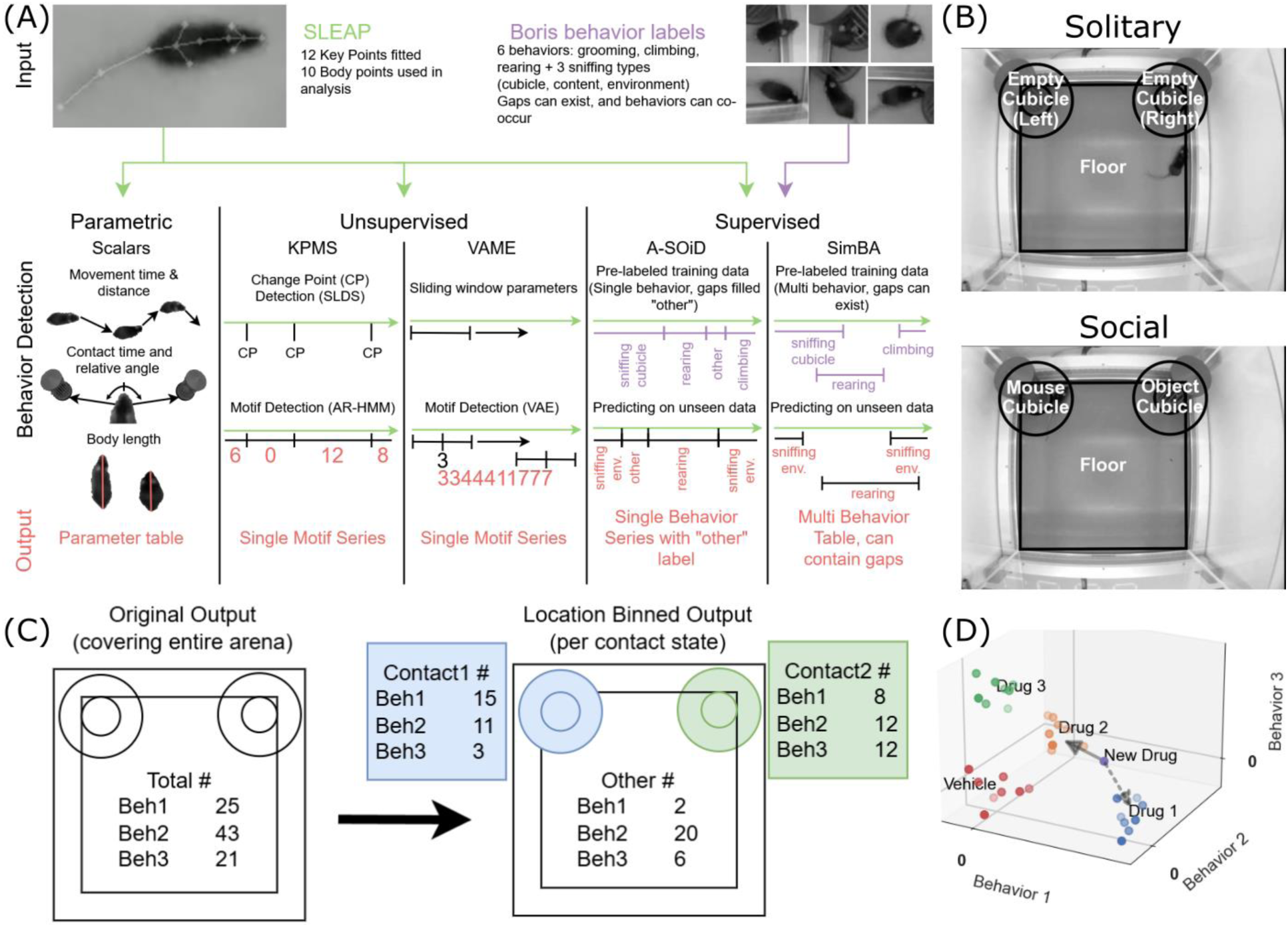
Experiment layout and parameters. (A) Input data (tracking and manual behavior labels) were fed into a selection of behavior summary models (e.g. Parametric, KPMS etc.). As a result of different architectures in each of these summary models, the final output also differs. Outputs can cover the entirety of the input time series, or only segments, and can include either a single behavioral label, multiple parallel labels, or a list of scalar parameters. (B) Top-down view of the arena, with relevant zones around the cubicles are overlaid in both “solitary” and “social” phase. (C) The outputs were further processed by grouping parameters and bouts in the zone they occurred in. Contacts were defined by a criterion of 100mm from a given zone as illustrated by blue and green colored circles, and the corresponding zones are shown in panel B. (D) The resulting features were normalized to the distribution of the pooled vehicle group. In the further analysis, models were evaluated by taking single samples and evaluating the closest treatment cluster in this feature space.

#### 2.6.1 Parametric behavior aggregation (Scalars)

As a baseline comparison we defined a set of classic scalar aggregations of the extracted key point data. The scalar parameters were selected with subject matter expert feedback based on experience from previous studies. Each phase recording was summarized as follows: the time and total distance spent moving, the time spent in contact with each cubicle, the relative angle to each cubicle (average and standard deviation), and the body length (average and standard deviation). These parameters were extracted from the entire dataset and then post-processed as described below.

We used the key point located at the base of the neck as a reference point for the position of the mouse, and the nose as the reference for the heading. We then converted these pixel-based measures to a distance-based measure using the pixel per millimeter resolution of our cameras, calculated based on the known distance between the corners of the arena and the same distance in pixel measured in the video (measure extracted through the Python package OpenCV2, version 4.11.0; using the `selectROI` function).

#### 2.6.2 Unsupervised behavior segmentation

The two tools, Keypoint-MoSeq (see below, Weinreb et al. 2024) and VAME (see below, Luxem et al. 2022), used for unsupervised behavior segmentation used the full set of key points tracked in SLEAP, except for two points located on the tail.

##### 2.6.2.1 Keypoint-MoSeq

We used Keypoint-MoSeq version 0.5.2 to extract behavioral “syllables” from the reduced SLEAP tracking dataset. The model was fitted to our vehicle training dataset, with a target median syllable length of 10 frames (∼400 ms). In a second step, the trained model was then applied to the full dataset.

Keypoint-MoSeq enables the detection of a threshold for filtering unstable key points. Through querying the user multiple times to relabel machine-labeled frames, it is possible to detect the level of detection “likelihood” below which the distance between machine-labels and ground truth manual labels increases dramatically. This likelihood threshold was detected in our dataset to be around ∼0.3945 (a.u.) and was also applied to the configurations of all other tools, if possible.

All downstream processing excluded syllables with a global onset proportion below 0.5%. This threshold was proposed in the analysis tools included in the Keypoint-MoSeq package and followed in this study. We named all 27 syllables with a bout proportion above 0.5% of the global distribution. For a full list and descriptions, see the Supplementary Methods.

##### 2.6.2.2 VAME

VAME version 0.10.0 was used to extract behavioral “motifs” from the key points extracted through SLEAP. We again truncated the key points to 10 by excluding the tail and modified the default parameters to fit with our existing data (see Supplementary Methods). Following the procedure established in literature, we first trained the model on the training data set and then segmented the same data into 100 modules. 41 of the found modules had an average bout proportion across all training recordings greater than 1%, and so this number of modules was applied for the segmentation of the entire dataset.

#### 2.6.3 Supervised behavior segmentation

We chose to use SimBA and A-SOiD as examples of supervised learning. Since the default SimBA feature set expects a specific selection of key points representing a subset of our key points, we filtered and renamed our base tracking dataset to match the same points (see Supplementary Methods).

To follow best practices, we split the BORIS-annotated training dataset in an 80:20 split into 30 videos to be used in the SimBA and A-SOiD training pipelines, and 6 validation videos to be held back and used to evaluate the trained models generated by both tools.

##### 2.6.3.1 SimBA

We used SimBA (Goodwin et al. 2024) Version 3.9.3 for supervised machine learning. During preprocessing, no outlier correction was performed, to ensure that the data used was the same as provided to the unsupervised methods. As the dataset was adapted to the default 8 keypoint SimBA format, we were able to use the default feature extraction pipeline. Six independent random forest classifiers were trained in a single step on the training dataset of 30 videos (see Extended Methods for configuration). Inference was run using a Jupyter notebook that applied the six trained models to the full dataset of 314 videos.

##### 2.6.3.2 A-SOiD

We used A-SOiD (Tillmann et al. 2024) version 0.3.1.3 as an example of active supervised learning. We followed the default pipeline (see Extended Methods for configuration) and trained the initial models on half of the training data (15/30 videos). The next 15 iterations were trained on 1 new video each, followed by retraining the models with the additionally labelled bouts. As the number of uncertain bouts shrunk in later iterations, we ran the last 3 iterations on the full iteration set of 15 videos, while labelling uncertain bouts in each video. We used the default threshold of 0.5 for the detection of uncertain bouts.

#### 2.6.4 Behavioral summary post-processing

All segmentation models returned a set of behavioral bouts of various frequencies and lengths (see Suppl. Figure 2D), including a start and stop frame index for each occurrence. This frame index corresponds to a video frame containing the location of the mouse within the arena. To capture location-dependent effects within the bout occurrences, we grouped occurrences into three possible states: contact with the enclosure containing the mouse, contact with the object enclosure, and no contact (Figure 2B, C). The contact criterion for either enclosure was a distance below 100mm, which we already established in our previous research (Ritter et al. 2025). Enclosures were named for their content during the “social” phase, e.g. the “mouse” enclosure during the solitary phase was the enclosure that contained a mouse in the social phase. For the parametric aggregation method, we calculated the scalar values per contact state for those values that allowed this procedure (body length, enclosure relative angles and distances, moved distances and time spent mobile, and total time in contact).

### 2.7 Sample classification and model evaluation

To evaluate the performance of all models, we analyzed their output on three levels, from here on called “tasks”: The detection of treatment effect compared to vehicle, the discrimination between treatments regardless of dosage, and the full discrimination of treatment and dosage pairs. Because of the varying dimensionality of the behavioral summaries, we chose to normalize the output of each summary model for all mice to the pooled vehicle control group and tested the classification of each data sample (behavior summary of a mouse) in a One-versus-Rest approach (Figure 2D), using nearest centroid classifiers based on the Manhattan distance. Specifically, for each task, we took a test data sample (i.e. behavioral summary of a mouse), then calculated the centroid of data samples in each of the treatment group and assigned the test data sample to a treatment group based on its Manhattan distance to the closest centroid. We repeated this approach for every data sample across all treatment groups in the three tasks. By following this procedure, it was possible to classify an unseen data sample (i.e., behavioral summary of a mouse) to one of the treatment groups within our dataset. In an ideal case, the models would assign each data sample to its corresponding treatment group (Figure 2D).

### 2.8 Statistical analysis

All statistical tests were performed in Python (version 3.11.9). Unless stated otherwise we used a significance threshold of α=0.05 and corrected for multiple testing by applying the Bonferroni correction where applicable.

#### 2.8.1 F-test

To determine which parameters showed significantly different variance between the vehicle group and a treatment-dosage group in Suppl. Figure 3A, we applied an F-test using the distribution implemented in Scipy (function `f.cdf`), with 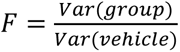 and *DOF* = *n_Samples_* − 1, and applied the Bonferroni correction for multiple testing.

#### 2.8.2 Manhattan distance

The Manhattan distance corresponds to the sum of differences across all dimensions between two points. In our case we apply the Manhattan distance to evaluate the distance between each sample and the centroids we consider in our analysis.

The equation for the Manhattan distance is 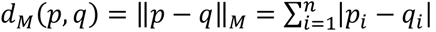, with p and q being two points in n-dimensional space.

#### 2.8.3 Closest centroid detection

To calculate the closest centroid for each sample we utilized the `NearestCentroid` class contained in the Scikit-Learn Python package, version 1.5.1. The prediction was done using cross-validation (function `cross_val_predict`) with the `LeaveOneOut` strategy, both contained in the same package.

#### 2.8.4 Binomial test and confidence interval

To test the recall of the models across detection tasks for significant differences, we modeled the outcome as a binomial result. Samples were either assigned correctly or not, with the chance level of the correct assignment based on the specific detection task:

- p_effect_=1/2 → correct treatment vs vehicle
- p_compound_=1/4 → correct compound vs 2 other compounds + vehicle
- p_treatment_=1/9 → correct treatment vs 7 other treatments + vehicle

The test statistic was calculated using SciPy’s binomial test function (function binomtest) and translated to a confidence interval using the normal distribution implementation (function norm).

We applied this test and interval in Figure 4B, Figure 5B, Figure 6B, Figure 7B, as well as Suppl. Figure 2E, Suppl. Figure 3C, Suppl. Figure 4C, Suppl. Figure 5B, and Suppl. Figure 7 to test for recall of various features (treatment, dosage, context) above chance level.

#### 2.8.5 Wilcoxon signed-rank test

We used the Wilcoxon signed-rank (WSR) test in the following analysis:

- In Figure 4B, Figure 5B, Figure 6B, and Suppl. Figure 7 we applied a two-sided test to test for significant differences of group-wise recall in solitary and social context
- To check for possible effects of our data modifications, we applied a two-sided test on the outcome of the modified datasets and the original dataset in Figure 7B, as well as Suppl. Figure 3C, Suppl. Figure 4C, and Suppl. Figure 5B

We used SciPy’s (version 1.14.0) implementation of the Wilcoxon signed-rank test (function wilcoxon) with significance levels set at α=0.05.

#### 2.8.6 Mann-Whitney U Test

We used the Mann Whitney U test (MWU) in the following analysis:

- Suppl. Figure 3B shows the outcome of applying a two-sided test to the recall of treatment groups with and without significant fingerprint features (across models), split by analytical level and context
- In Suppl. Figure 6A we applied a two-sided test to compare performance between solitary and social context for a given model and analytical level. In addition, we also compared performance between different models within the same context and analytical level.
- To verify that within- and between-cluster distances differed, we applied a two-sided test in Suppl. Figure 6B to the distances yielded by each model

We used SciPy’s (version 1.14.0) implementation of the Mann-Whitney U Test (function mannwhitneyu) with significance levels set at α=0.05.

#### 2.8.7 Two-way ANOVA

We used the `ols` class and `stats.anova_lm` function implemented in the statsmodels library (version 0.14.2) to apply a two-way type 2 ANOVA to the global recall shown in Figures 4-6B. The applied formula was “`recall ∼ C(segmenter) * C(phase)`”.

#### 2.8.8 Bonferroni correction

All statistical tests that were applied to more than one comparison were corrected for false-positive discovery by applying the Bonferroni correction. Unless stated otherwise we used the significance threshold of α=0.05 and corrected according to **α_Bonferroni_** = **α_uncorrected_**⁄**n_tests_**, with **n_tests_** corresponding to the number of tests performed. This correction was applied to the binomial tests (**n_tests_** = 10) and WSR tests (**n_tests_** = 5) in Figures 4-6B; the binomial tests (**n_tests_** = 90) and WSR test (**n_tests_** = 60) in Figure 7B; the binomial tests shown in Suppl. Figure 2E (**n_tests_** = 5); the F-test in Suppl. Figure 3A (**n_tests_** = 13500); the MWU test in Suppl. Figure 3B (**n_tests_** = 3); the binomial tests (**n_tests_** = 60) and WSR tests (**n_tests_** = 30) in Suppl. Figure 3C; the binomial tests (**n_tests_** = 66) and WSR tests (**n_tests_** = 36) in Suppl. Figure 4C; the binomial tests (**n_tests_** = 48) and WSR tests (**n_tests_** = 18) in Suppl. Figure 5B; the MWU tests between contexts (**n_tests_** = 15) and between models (**n_tests_** = 60) in Suppl. Figure 6A; the MWU tests (**n_tests_** = 10) in Suppl. Figure 6B; the binomial tests (**n_tests_** = 100) and WSR tests (**n_tests_** = 50) in Suppl. Figure 7.

## 3 Results

We recorded videos of 157 male C57BL/6J mice in an open field test (OFT) with two phases. Each mouse was injected with a compound or vehicle before the start of the recording (see Methods) and placed in the arena after a delay of 20 minutes. During the first, “solitary” recording phase (10 minutes) the mice were alone in the open arena, except for two empty cubicles.

**Figure 3.**
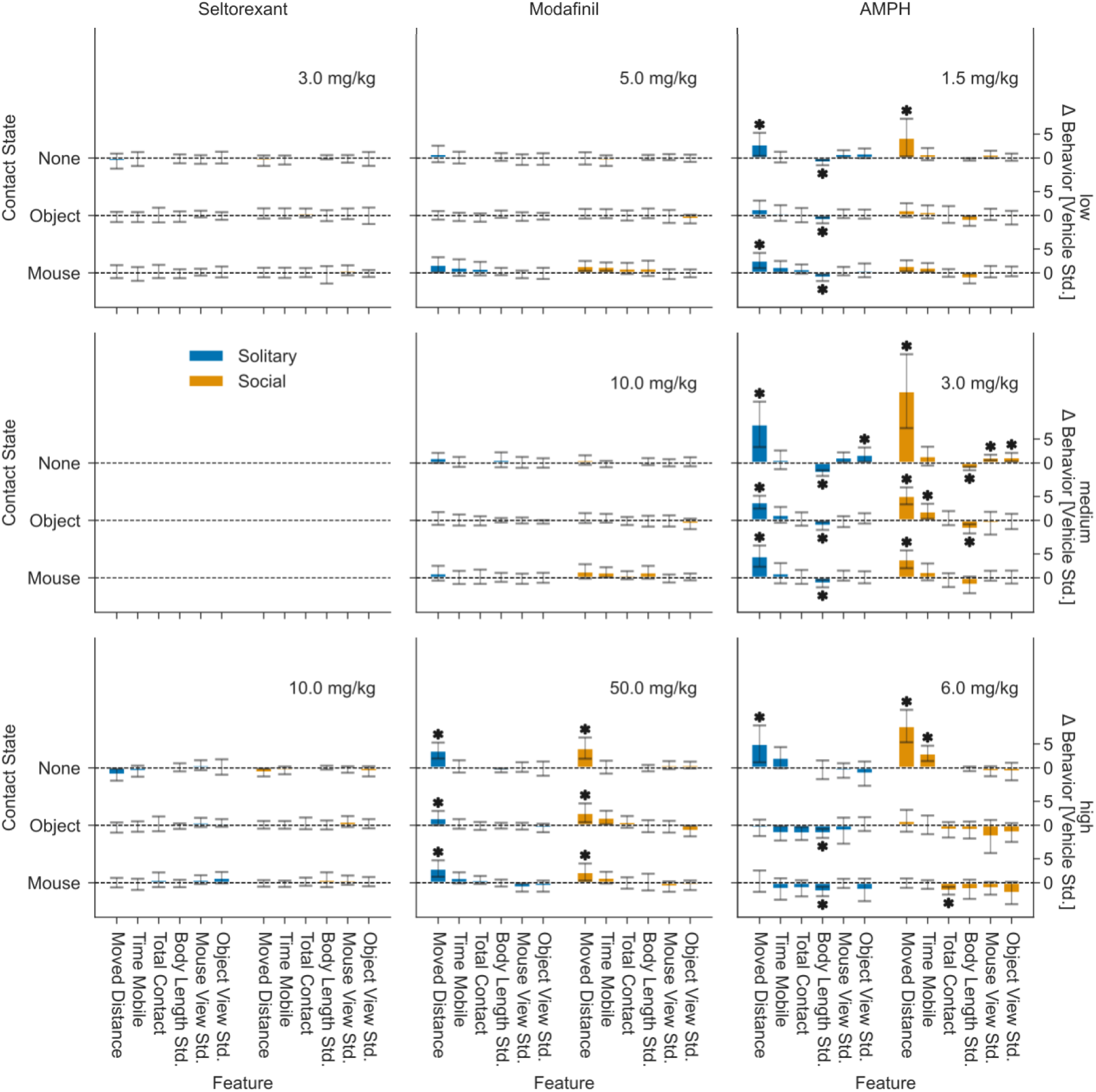
Comparison of treatment effects across locomotion parameters. Bar plots showing vehicle-relative mean and standard deviation of individual locomotion parameters across treatments (columns), dosages (rows), and contact states. Y-axis on the left shows categorical contact states and y-axis on the right shows the quantification for the corresponding parameter. Significant differences marked with an asterisk (two-sided Mann-Whitney U Test, Bonferroni Correction, p<=0.05).

During a short break, an untreated second mouse from a different cage and a novel object were placed in the two corresponding cubicles respectively. The break was followed by the second “social” phase (10 minutes). Placement of object and mouse in the cubicles was counter-balanced across recordings and later normalized to the mouse being in the left enclosure during analysis. All mice were housed under normal light cycle conditions in the same facility. Experiments were performed during the light phase. Using SLEAP (Pereira et al. 2022) we tracked 12 points across the body of the freely moving mice. This source dataset was then filtered to a subset of 10 body points for parametric and unsupervised behavior analysis, and 8 points for supervised behavior analysis (see Methods).

First, to verify findings of our previous study, we tested the ability of different behavioral summary models to detect context within recordings of vehicle treated mice (see Suppl. Figure 2E; Binomial test, Bonferroni correction, p<=0.05). Our statistical tests showed context separation significantly above chance for all models, indicating that context-dependent differences in behavioral phenotype are recognizable across models. This also aligns with the findings of our previous study, which we were able to reproduce in this vehicle-treated control group (Ritter et al. 2025).

Before evaluating behavior summary models on detection of treatment effect and discrimination of drug-dose combinations, we verified the treatment effects on the locomotion using 6 different parameters calculated from the key point data (Figure 3; see Methods). Analysis of changes in these 6 parameters in comparison to the pooled vehicle treatments showed that treatments elicited the expected behavioral outcomes across ‘solitary’ and ‘social’ context (Figure 3). We observed little effect of seltorexant (left column in Figure 3), even at high dosage, but were able to identify significant increases in moved distance in high dosages of modafinil (middle column in Figure 3) and amphetamine (right column in Figure 3; Mann-Whitney U Test, Bonferroni Correction, p<=0.05). It is noteworthy that this effect was only visible for high-dose amphetamine when mice were not in contact with one of the enclosures, independent of context (bottom right subplot in Figure 3). During contact with either enclosure, amphetamine-treated animals showed a significant reduction in the standard deviation of the body length measurement in solitary context (right column in Figure 3). Additionally, high dosage amphetamine-treated animals spent more time mobile without contacts and spend less time with the conspecific in social context. In comparison, medium dosage amphetamine treatment led to a significant increase of moved distance independent of contact or context, as well as a reduction in body length standard deviation in all cases except for contacts with conspecifics.

### 3.1 Treatment effect detection is similar across behavioral summary models

To evaluate the impact of different summary models in detecting an effect of a drug treatment, we compared each treatment-dosage group to the pooled vehicle group. Each model was evaluated based on the assignment of a given mice to a treatment group or pooled vehicle group using its behavioral summary measures (see Methods). Recall was calculated as the number of correctly assigned samples over the total number of samples.

With no exceptions, the models were able to detect effects across treatments independent of context (Figure 4A). Further analysis showed that all models performed above chance (Figure 4B; two-sided binomial test, Bonferroni correction, p≤0.05), with none of the pairwise comparisons between solitary and social context being significant (two-sided Wilcoxon signed-rank test, Bonferroni correction, p≤0.05).

**Figure 4.**
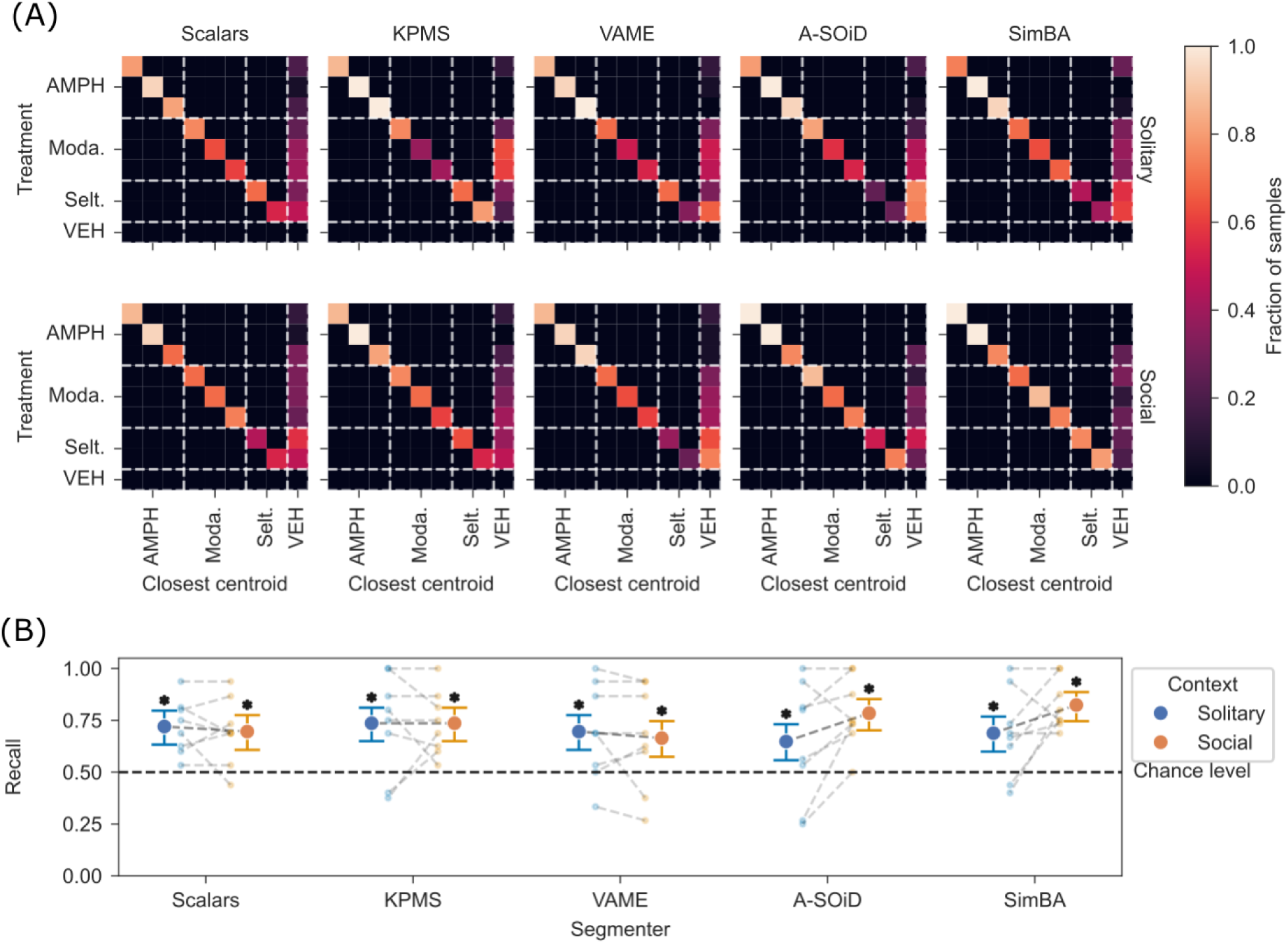
Quantification of treatment effect detection across models and contexts. (A) Confusion matrices in a 5 x 2 grid, representing the performance of the five models in the solitary and social contexts. Rows are actual treatments, while columns show the treatment assigned by the nearest-centroid classifier. Color scale represents the fraction of samples in each assignment. Only assignments on the diagonal (identical treatment and classification) are correct. Misassignments are shown outside the diagonal. (B) Point plots (CI=95%) show average recall for each summary model in solitary and social context. Recall values for each compound treatment in both contexts are shown in the background (dotted lines). Recall values significantly (two-sided binomial test, Bonferroni correction, p≤0.05) above chance level are marked with an asterisk. Differences between contexts were also tested (two-sided Wilcoxon signed-rank test, Bonferroni correction, p≤0.05) but did not result in significance.

A two-way ANOVA (see Methods; Bonferroni correction) revealed no significant main effects or interactions for behavior summary models and contexts. A deeper look into each models’ behavioral summary suggested that the number of parameters involved in the “fingerprint” of the individual treatment-dosage effects varied widely across models (see Suppl. Figure 3A; F-statistic, Bonferroni correction, p≤0.05), independent of the type of model. These results show that the mere detection of a treatment effect is possible across established models and methods and is not affected by the choice of context in this study.

### 3.2 Treatment discrimination performance is similar across behavioral summary models

As there was no significant difference in performance across models and contexts when limiting analysis to detect treatment effects, we investigated the models’ ability to discriminate between drug treatments regardless of dosage. For that purpose, we again classified all mice into treatment and vehicle groups with a nearest-centroid classifier (see Methods). If the closest centroid was of the correct treatment group (or vehicle) independent of dosage, the assignment was classified as correct. All models were able to correctly assign samples to their treatment, but the overall performance was slightly reduced (see Figure 5A) when compared to the detection performance (Figure 4). Applying the same tests as before showed that all models performed significantly above chance level (see Figure 5B; two-sided binomial test, Bonferroni correction, p≤0.05), with no significant effect of context (two-sided Wilcoxon signed-rank test, Bonferroni correction, p≤0.05). A second two-way ANOVA with behavior summary models and contexts as factors revealed neither significant main effects, nor significant interactions. Unexpectedly, these results suggest that there is neither a dependence on a certain method or model to discriminate between the effects of treatments, nor a significant influence of context.

**Figure 5.**
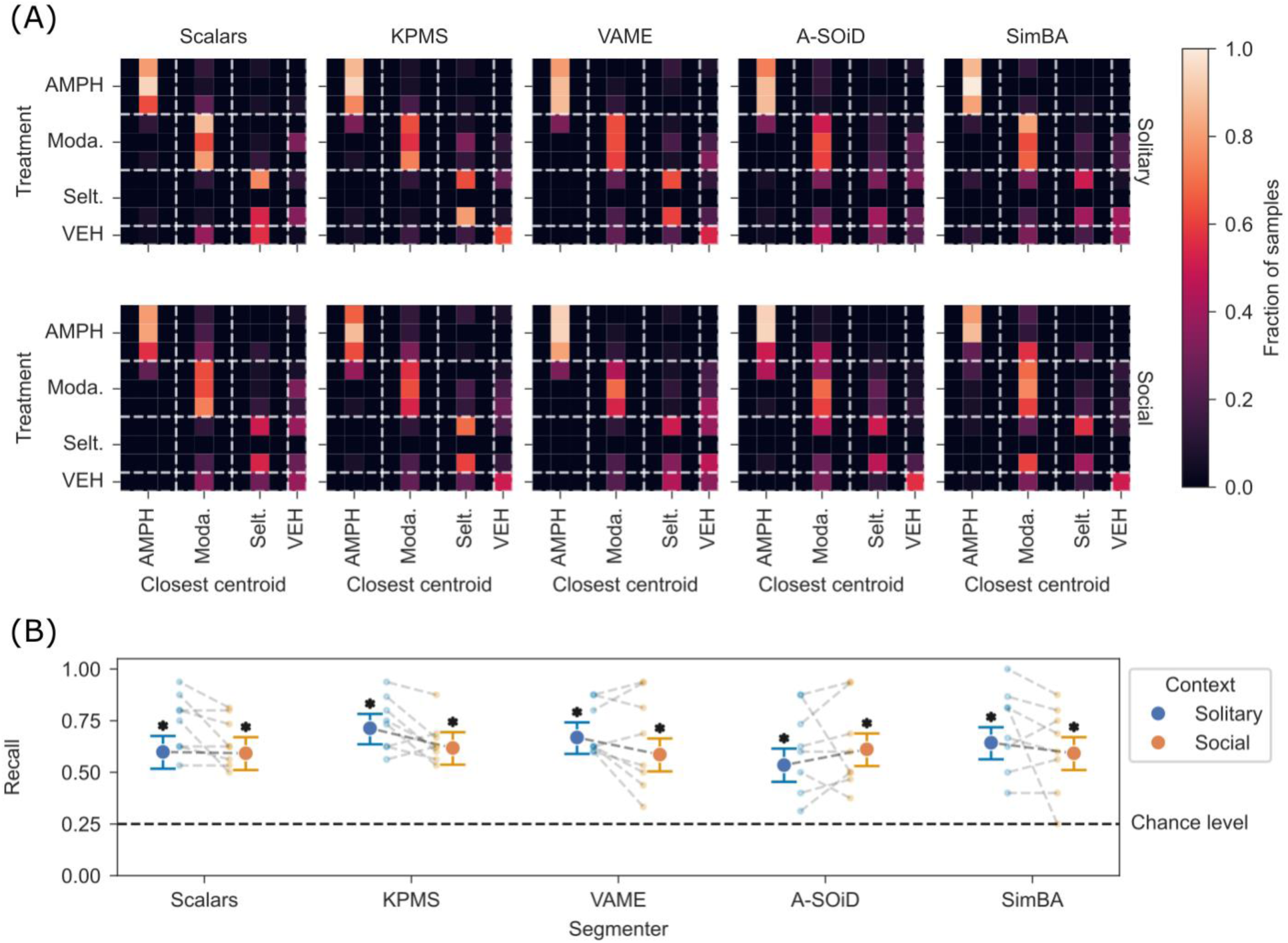
Quantification of treatment discrimination across models and contexts. (A) Confusion matrix for treatment discrimination, same layout as in Figure 4A. (B) Point plots for treatment discrimination, same layout and tests as in Figure 4B.

**Figure 6.**
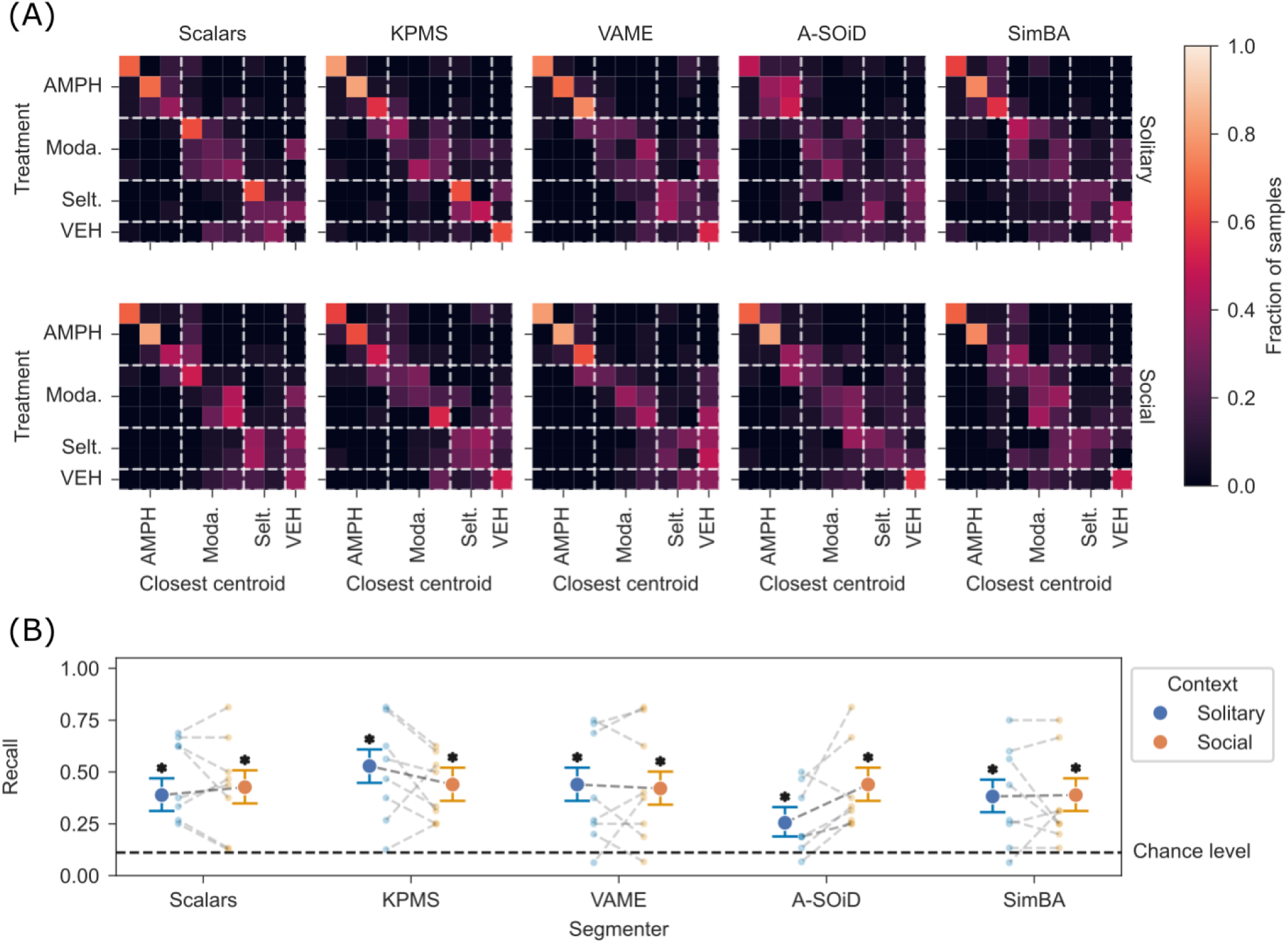
Quantification of treatment-dosage discrimination across models and contexts. (A) Confusion matrix for treatment-dosage discrimination, same layout as in Figure 4A. (B) Point plots for treatment-dosage discrimination, same layout and tests as in Figure 4B.

### 3.3 Treatment-dosage discrimination performance is similar across behavioral summary models

Since there were no significant differences in performance when trying to discriminate between treatments, we tested the summary models’ ability to separate full treatment-dosage pairs in our dataset. Each sample was mapped to its closest centroid, but now only the assignment to the correct treatment as well as dosage was classified as a correct prediction. The performance of all models in this task was strongly degraded compared to the two previous tasks, except for the separation of the amphetamine treatment-dosage groups (see Figure 6A). With the same statistical approaches as described before, we were able to show that all models performed above chance, with no significant effect of context on each model’s performance (see Figure 6B; two-sided binomial test & Wilcoxon signed-rank test, Bonferroni correction, p≤0.05). Applying ANOVA showed again no significant main effects or interactions. This surprising result suggests that in a detailed comparison of effects across different treatments and dosages, the performance of parametric, supervised, and unsupervised behavior summary models is not significantly different.

### 3.4 Contribution of contextual features to detection and discrimination of treatments

Finally, we tested whether there were distinct contextual features that the summary models captured to achieve the observed detection and discrimination of treatments.

**Figure 7.**
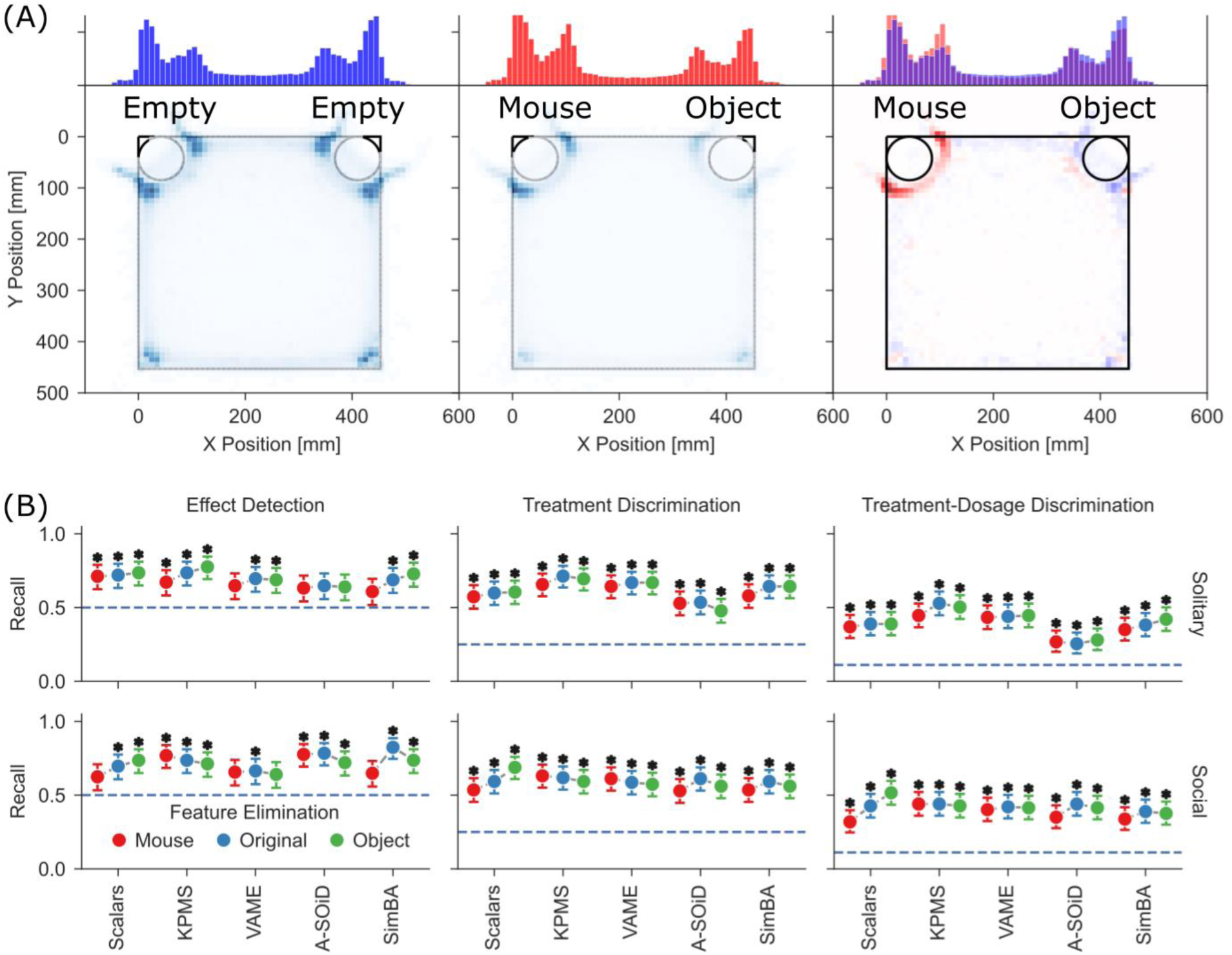
Quantification of treatment detection and discrimination with elimination of contextual features. (A) Mouse location histograms for solitary context (left), social context (middle), and the difference between social and solitary context (right). Upper plots show only binned x-position, lower show both dimensions. (B) Average recall across treatments for different behavioral summary models in solitary (top row) and social contexts (bottom row) with elimination of features associated with mouse or object cubicles in comparison no feature elimination. Same statistical tests and annotation as in Figures 4-6.

A recent study (Wiltschko et al. 2020) showed that only a characteristic subset of features is required to discriminate treatments. We explored this aspect by applying an F-statistic to detect features that could be used as a “fingerprint” of a particular treatment compared to the control group. We found that the presence of “fingerprint” features in a given treatment sample resulted in significantly increased detection and treatment discrimination performance, but only during solitary context (see Suppl. Figure 3B; Mann-Whitney U Test, p<=0.05). Yet, eliminating the “fingerprint” features and repeating the same analysis as shown in Figures 3-5B did not change the outcome significantly, suggesting that these “fingerprint” features in a summary model do not contribute significantly to above chance detection and discrimination (see Suppl. Figure 3C; Wilcoxon signed rank test, Bonferroni Correction, p<=0.05).

We observed that position preference of the freely moving mice was strongly affected by presence of the second mouse in the cubicle during the recording of the social context (see Figure 7A). To test whether there were features particular to contact with a mouse or object enclosure that contributed to the treatment detection and discrimination, we dropped features related to contact with either mouse enclosure or object enclosure. While eliminating mouse enclosure contact features in social context recordings led to some of the models, for example parametric aggregation, to stop performing significantly above chance level. However, none of the effects were significantly different to the performance on the original dataset (see Figure 7B). Similarly, eliminating features related to object enclosure contact did not lead to significant changes.

We observed that behavior summary from the supervised models (e.g. SimBA) did not cover the full recording unlike the unsupervised models (e.g. VAME). To address if this difference between the models played a role in detection and discrimination, we limited the temporal occurrence of the behavior summary of unsupervised models to that of supervised models and repeated the analysis. Surprisingly, results did not lead to significant changes in performance (see Suppl. Figure 4). Furthermore, extension of the feature vectors to include the transition probabilities between behavioral motifs, in relevant models (KPMS, VAME, A-SOiD), did not lead to significant changes in performance (see Suppl. Figure 5).

To investigate whether there was a minimal set of features per model that allowed performance significantly above chance levels within a particular context, we repeated the analysis for every feature separately. There were 536 possible model/context/behavioral motif combinations. Filtering combinations that did not occur for the vehicle control group to avoid overestimating performance led to a total of 29 combinations being removed. The remaining combinations were re-analyzed in all three detection and discrimination tasks, leading to a total of 307 occurrences of single features allowing significantly above chance performance (∼20% of all possible tests, p<=0.05, binomial test, Bonferroni correction). The feature set for p<=10e-5 (chosen for visualization purposes) along with the respective global recall can be found in Suppl. Table 1.

To further verify the outcome of this study, we extended Figures 4-6B by comparing the relative distance (separation) between the correct group centroid, and the closest incorrect centroid. This allowed us to estimate performance beyond the mere correct or incorrect assignment summarized in the recall metric. Supervised models separate significantly better in social context than VAME for effect detection (Mann Whitney U Test, Bonferroni Correction, p<=0.05), and for treatment-dosage discrimination unsupervised models show better separation than A-SOiD in solitary context (see Suppl. Figure 6A; Mann Whitney U Test, Bonferroni Correction, p<=0.05), but A-SOiD significantly improved separation in social context.

Overall, these results show that the tested common behavioral summary methods are usually and reliably sufficient for detection and discrimination of treatment effects at a level significantly above chance, at least in the treatments and dosages chosen in this study.

## 4 Discussion and conclusion

The increasing number of methods available to summarize complex behaviors of mice makes the optimal choice of a model for a behavioral paradigm (e.g. social or solitary) and drug treatment detection and discrimination more difficult. However, there has not been an attempt to evaluate performance of these models across behavioral context and levels of treatment detection and discrimination. In this study, we recorded freely moving mice during exploration of an open field arena in solitary and social contexts (see Figure 1) after being treated with 8 different treatment-dosage pairs (2 seltorexant, 3 modafinil, 3 amphetamine; see Methods). Next, we applied five established behavioral summary models: parametric aggregation (see Figure 2A), KPMS (Weinreb et al. 2024), VAME (Luxem et al. 2022), A-SOiD (Tillmann et al. 2024), and SimBA (Goodwin et al. 2024), to evaluate treatment detection and discrimination of these models based on their summary outputs. Surprisingly, we found that the choice of model did not have a significant effect on the detection of a drug treatment effect relative to vehicle (Figure 4), discrimination between drug treatments regardless of dosage (Figure 5), and discrimination between drug-dosages (Figure 6).

While the input and output of the models differed (see Suppl. Figure 2C), their performance remained significantly above chance level for all tested discrimination levels, with no statistically significant differences between the tested model and context combinations. Selection of specific summary features (e.g. mouse enclosure contacts features) also yielded no consistent and significant changes in recall (see Figure 7B). We also addressed the potential issue of the behavioral label coverage that differed widely between models and contexts, by limiting higher label coverage to a subset of frames or by eliminating “unsupervised-specific” motifs from the summary output, and the results did not significantly affect performance (see Suppl. Figure 4C). Furthermore, we also tested if transitions between behavioral motifs played a significant part in detection and discrimination of treatments (see Suppl. Figure 5A), and the results showed no significant changes in recall (see Suppl. Figure 5B).

We also considered if limitations to (analog) frame rate and training set (see Methods) may have played a role in the model extraction of behavioral summary to detect and discriminate treatments. We argue that these limitations are the same across all models, and all models perform significantly above chance level, across all treatment detection and discrimination levels, and can separate context (see Suppl. Figure 2D).

Further limitations of our study may include the use of relatively low number of mice and the limited set of treatments compared to previous studies (Wiltschko et al. 2020). As the number of mice correlated with the number of treatment-dosage pairs, we compared groups of 16 to 32 mice, group sizes that are in line with established literature and our prior research. Finally, we selected the treatments and dosages shown in this study based on previous literature (Simon et al. 1996; Bonaventure et al. 2015; Yates et al. 2007). All three treatments (amphetamine, modafinil, and seltorexant) have been previously used by our lab, and the discrimination between amphetamine and modafinil has been shown previously to be potentially challenging (Wiltschko et al. 2020) and was of interest in this study. Outcomes between seltorexant (reduced activity, increased sociability) and the other two treatments (increased activity) was expected to have a clear contrast (Muehlan et al. 2020). Thus, this small set of three treatments is well chosen to compare the discriminative ability of behavioral summary models. Similarly, another potential limitation of the paradigm may be the role of enclosures for stationary second mice and the novel object. We have addressed this aspect in our previous study (Ritter et al. 2025) and our results showed that the having second mice in an enclosure lead to the same general effects of social context as a second freely moving mice (see also Suppl. Figure 2D).

While the number of mice was in line with existing research, and appropriate for the number of treatments under investigation, the application of unsupervised segmentation methods leads to a high number of output parameters (behavioral motifs). We applied cross-validation, classification, and analysis methods that are independent of the number of parameters (Nearest Centroid classification using Manhattan distance with Leave-One-Out cross validation) and the outcome of our analysis is independent of classifier choice (see Suppl. Figure 7). This choice of not applying different classification models on the summary output, other than nearest centroid, was to allow us to evaluate the summary models in their own pharmaco-behavioral space.

To align our research with analysis shown in previous literature, we applied a comparison featured in Wiltschko et al. (2020), and calculated the within-group and between-group distribution of Manhattan distances across models and contexts (see Suppl. Figure 6B). Comparisons of within-group and between-group distances showed significantly larger between-group distances than within-group distances for all comparisons (Mann-Whitney U Test, Bonferroni Correction, p<=0.05). We then calculated the sample-wise ratio of the average between-group distance over within-group distance (see Suppl. Figure 6C). The distance ratios in our dataset show a similar range across models and contexts, with basic parametric aggregation showing a better ratio in social context when compared to other models. However, this outcome does not translate to better recall in our previous analysis. When applying a similar ratio calculation to the values shown by Wiltschko et al. (2020), it appears that the resulting values do not disagree with our findings, with both scalar aggregation and Keypoint-Moseq leading to similar between-group / within-group distance ratios. Furthermore, two out of three treatments (modafinil and amphetamine) selected in this study have also been shown in the same publication to have a low degree of separation. In addition, we applied the two classifiers also featured in their study, Logistic Regression and Support Vector classifiers, and the results aligned with our findings described above (see Suppl. Figure 7, rightmost column).

In summary, while the scope of our study is more focused on a small, but diverse set of treatments, we applied a wide range of models across two very different behavioral contexts. Building on our previous study (Ritter et al. 2025), we expected a significant influence of the presence of a conspecific on the treatment-related behaviors, and thus the ability to discriminate between them. Surprisingly, our results show that neither model choice nor context led to significantly different recall in our set of treatments. While the higher-dimensional output gained from unsupervised machine learning approaches leads to higher absolute distances between samples in their feature space, this does not result in greater relative separation of treatments. In the same way, there are significant changes in behavior in the presence of a conspecific, detectable with all tested models, yet this effect does not improve treatment recall. While this outcome does not contradict previous research, it also suggests that further research with an extended treatment set is needed, as it is possible that within the scope of this study the chosen treatments “saturate” their effects already in solitary conditions. On the other hand, it could be necessary to analyze both social behavior and treatment effects within a more complex, semi-naturalistic setting (Forkosh et al. 2019) to unlock possible interactions between both. Finally, our study suggests that the choice of models and behavioral context in pre-clinical research should be informed not just by performance metrics, but also by explainability, computational costs, and the specific research questions and treatments used in each study.

## Table of abbreviations

| Abbreviation | Full Term |
| --- | --- |
| KPMS | Keypoint-MoSeq |
| VAME | Variational Animal Motion Encoding |
| SimBA | Simple Behavioral Analysis |
| B-SOiD | Behavioral Segmentation of Open field in DeepLabCut |
| LED | Light Emitting Diode |
| OFT | Open-Field Test |
| PO | Per os, orally |
| SC | Subcutaneously |
| IP | Intraperitoneally |
| MWU | Mann Whitney U Test |
| WSR | Wilcoxon Signed-Rank Test |

## Acknowledgements

We want to thank the other members of the data science team for their feedback on this project.

## Data availability statement

The original contributions presented in the study are publicly available. This data can be found here: https://doi.org/10.5281/zenodo.21460909

## Code availability statement

The code used to generate the data shown in this study is publicly available. It can be found here: https://github.com/Marti-Ritter/comparison-multiple-video-tracking-based-behavioral-summary-approaches-for-compound-discrimination

## Ethics statement

All procedures followed the regulations for animal experimentation enforced by the local district administration’s animal welfare commissioner of the state of Baden-Württemberg.

## Author Contributions

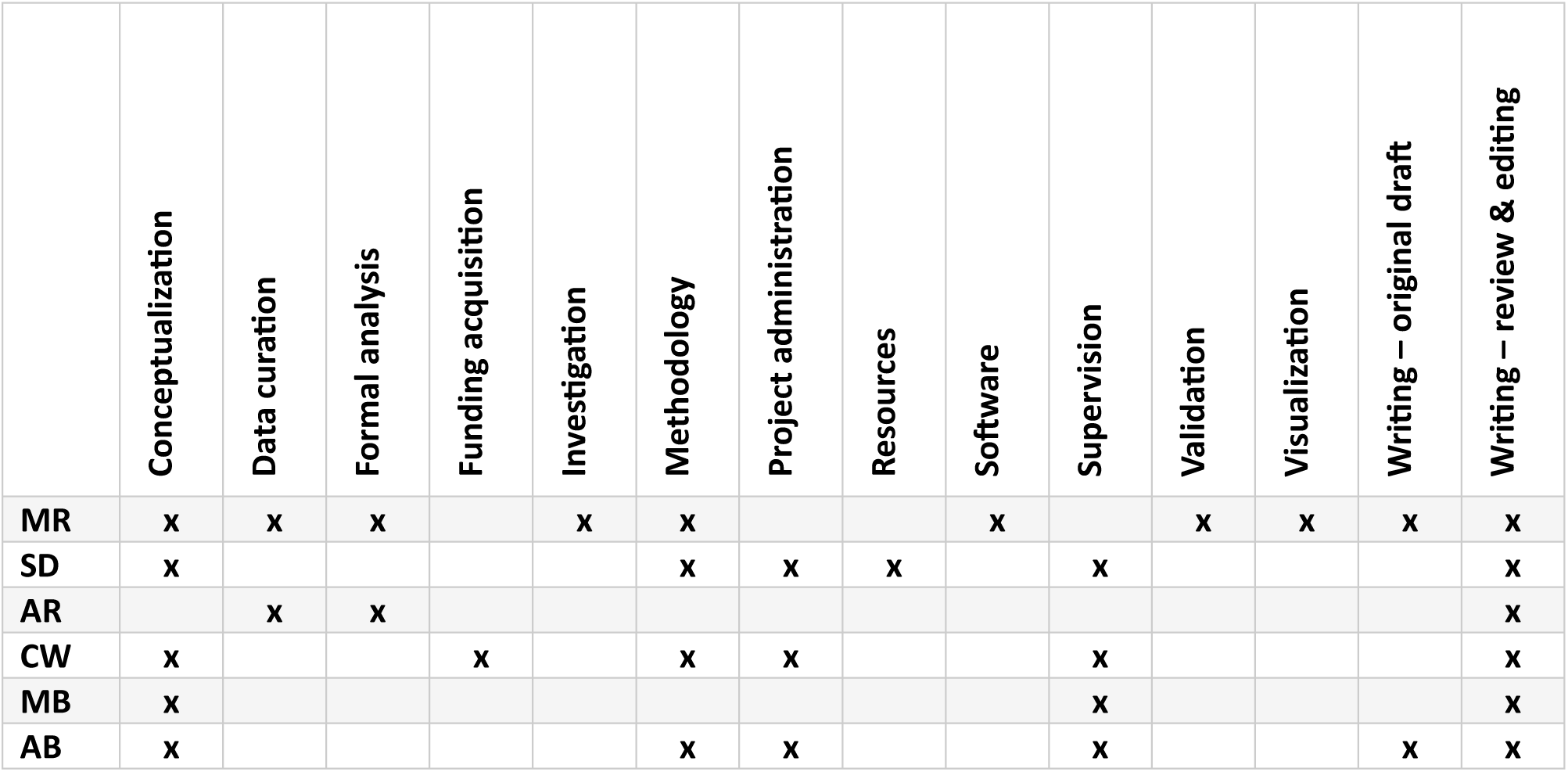

## Generative AI statement

The authors declare that no Generative AI was used in the creation of this manuscript.

