## Supplements for "Comparison of multiple video tracking-based behavioral summary approaches for compound discrimination"

### 1 Supplements

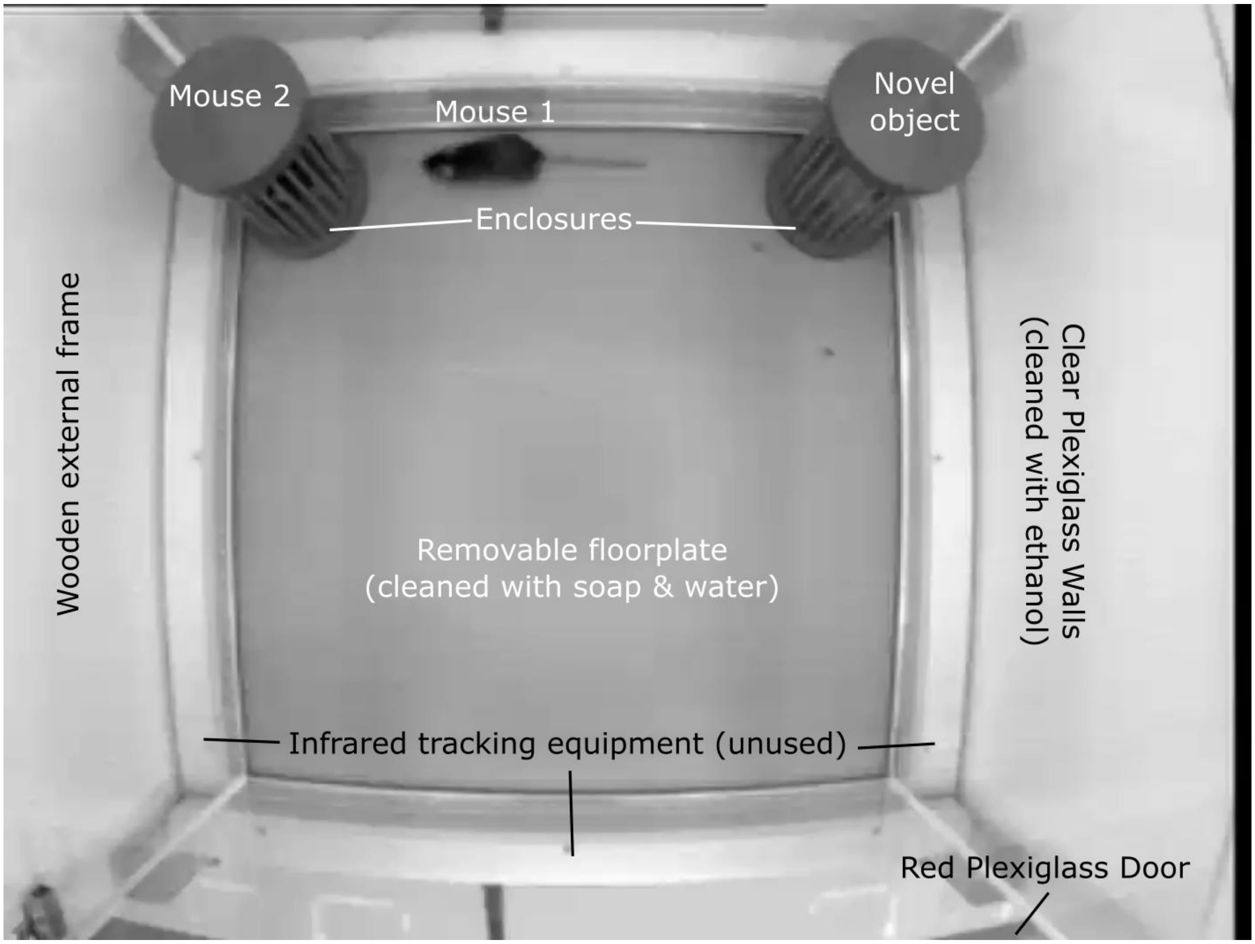


**Suppl. Figure 1. Top-down view of the setup.** In both solitary and social context, the shown setup was used. The walls were made of clear plexiglass, with the removable floor made of gray plastic. Between individual recordings the walls were cleaned with ethanol, and the floor with soap and water. The front (seen at the bottom of the image) was covered by a red plexiglass door. Around the perimeter of the floor infrared tracking equipment is visible, which was unused in this study. One mouse can be seen freely moving between two enclosures. In the solitary context both enclosures were empty. This frame is taken from a social context recording and a second mouse and a novel object can be seen in the two cubicles. Cubicle content was counterbalanced (swapped) across recordings and within groups.

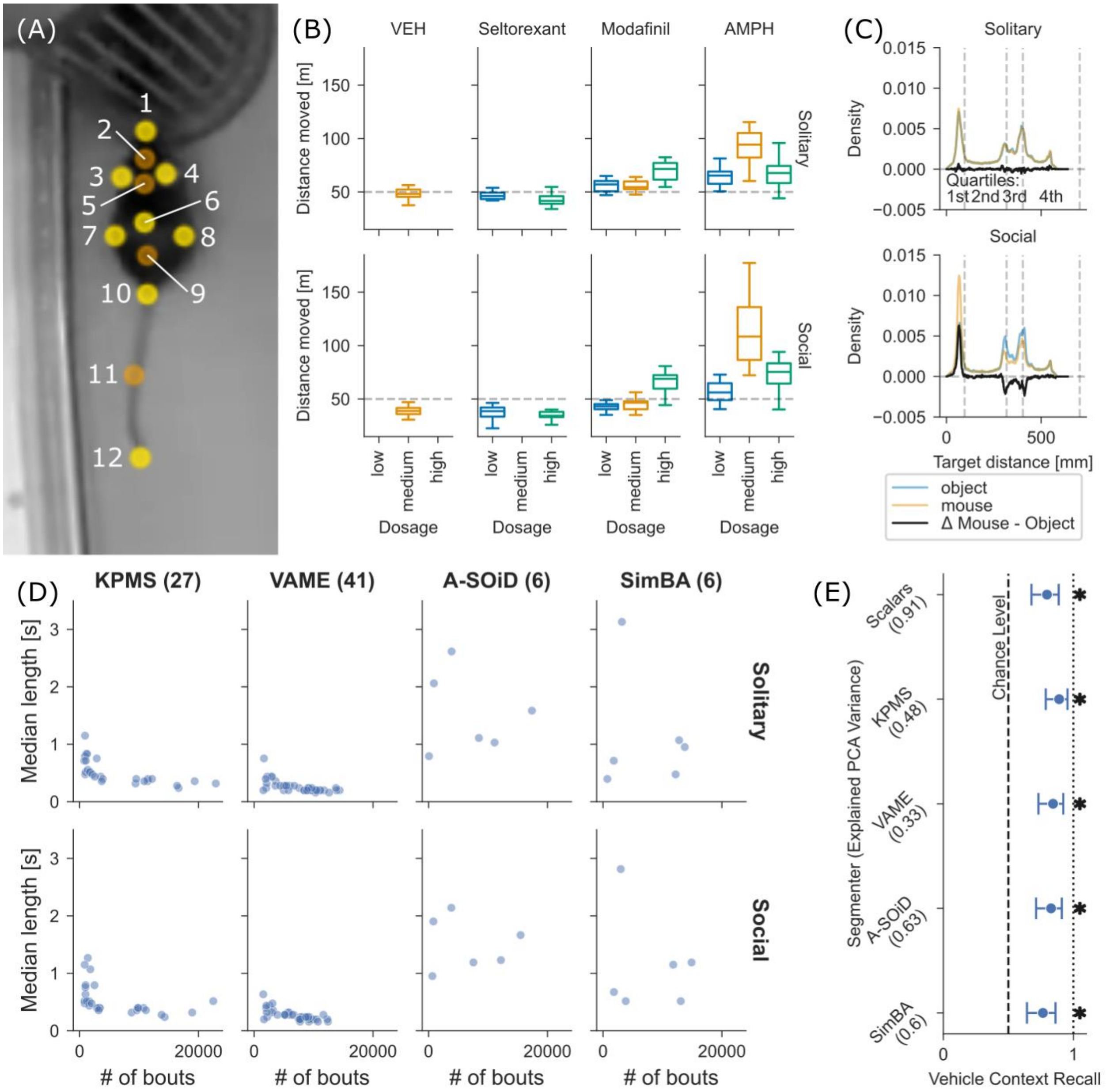


3

**Suppl. Figure 2. Overview over available data and extracted behaviors.** (A) Tracked keypoints. Keypoints that were dropped in the reduced SimBA set in orange. For names assigned to points see Supplementary Methods. (B) Boxplots of moved distances in the 9 treatment/vehicle groups. Dashed line represents solitary vehicle mean distance to facilitate comparison of distances. (C) Histogram of enclosure-relative distances in solitary and social context. Black line represents histogram difference between mouse and object enclosure. (D) Scatterplot showing relationship between frequency of behaviors and the lengths of their bouts. Number of dots in each facet correspond to the number of behaviors detected by each segmenter. (E) Global recall of context in vehicle animals. All comparisons were significantly above chance (binomial test, Bonferroni correction, p<= 0.05).

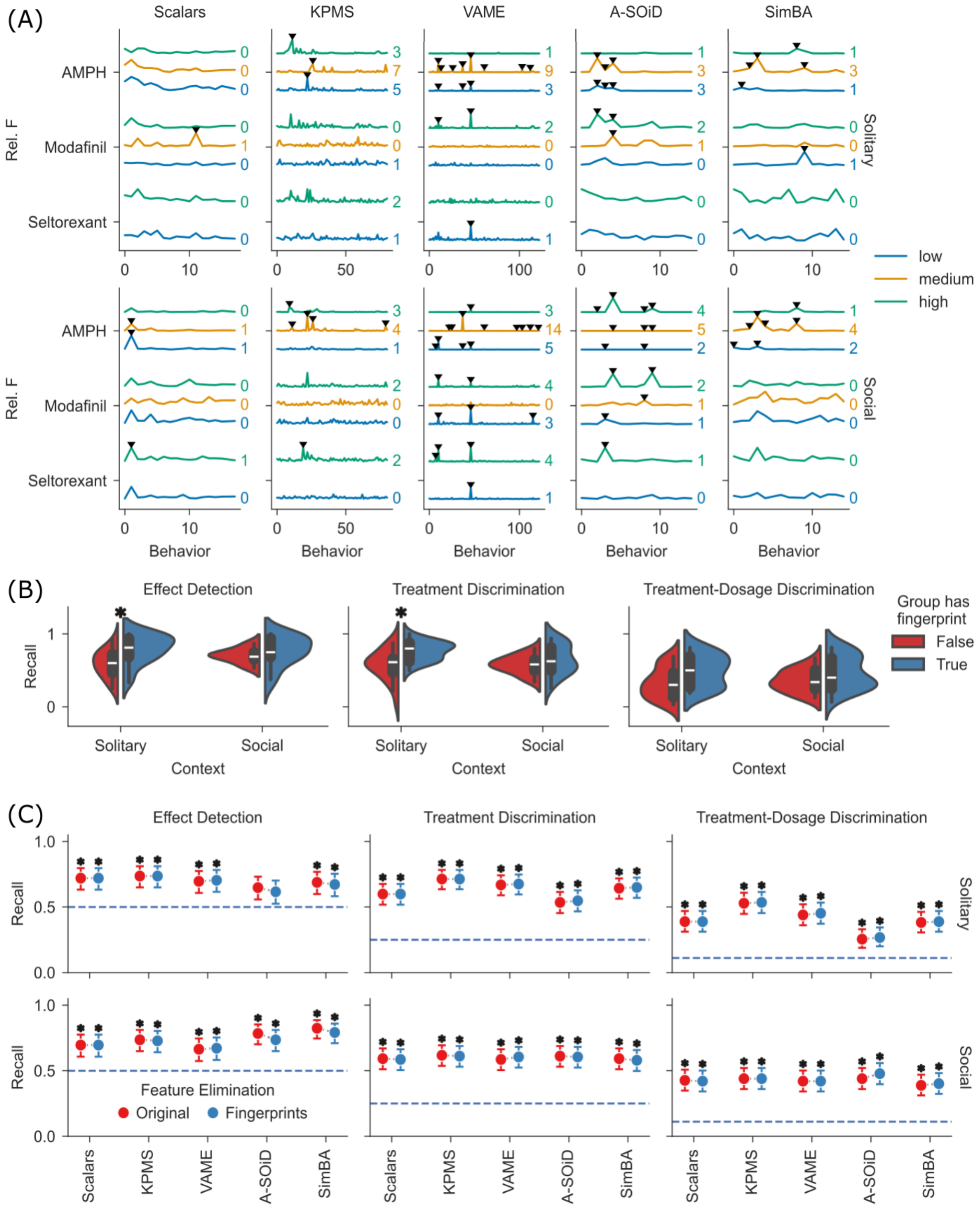

(A)
Scalars
KPMS
VAME
A-SOiD
SimBA
AMPH
Modafinil
Seltorexant
Rel. F
Solitary
Social
Behavior
low
medium
high
(B)
Effect Detection
Treatment Discrimination
Treatment-Dosage Discrimination
Recall
Solitary
Social
Context
Group has fingerprint
False
True
(C)
Effect Detection
Treatment Discrimination
Treatment-Dosage Discrimination
Recall
Feature Elimination
Original
Fingerprints
Scalars
KPMS
VAME
A-SOiD
SimBA
Solitary
Social


4

**Suppl. Figure 3. Overview of behavioral fingerprints and result of truncating them.** (A) Behavioral F-statistic fingerprint of each treatment. Columns correspond to individual segmenters, rows to context. Statistically significant behaviors are marked with a caret. (B) Violinplots of group recall across tasks. Plots are split between groups that have fingerprint features (see panel A), and those that do not. Significant comparisons between both are marked with asterisks (Mann-Whitney U Test, Bonferroni Correction, $p<=0.05$). (C) Global recall across tasks, context, and behavioral summary models. Input data to the classification was modified by dropping any features that were detected as significant per group. The outcome is compared against the unmodified dataset. Same tests and annotation as in Figures 4-6. Difference in significance in the original dataset is caused by different numbers of multiple tests being performed and corrected for.

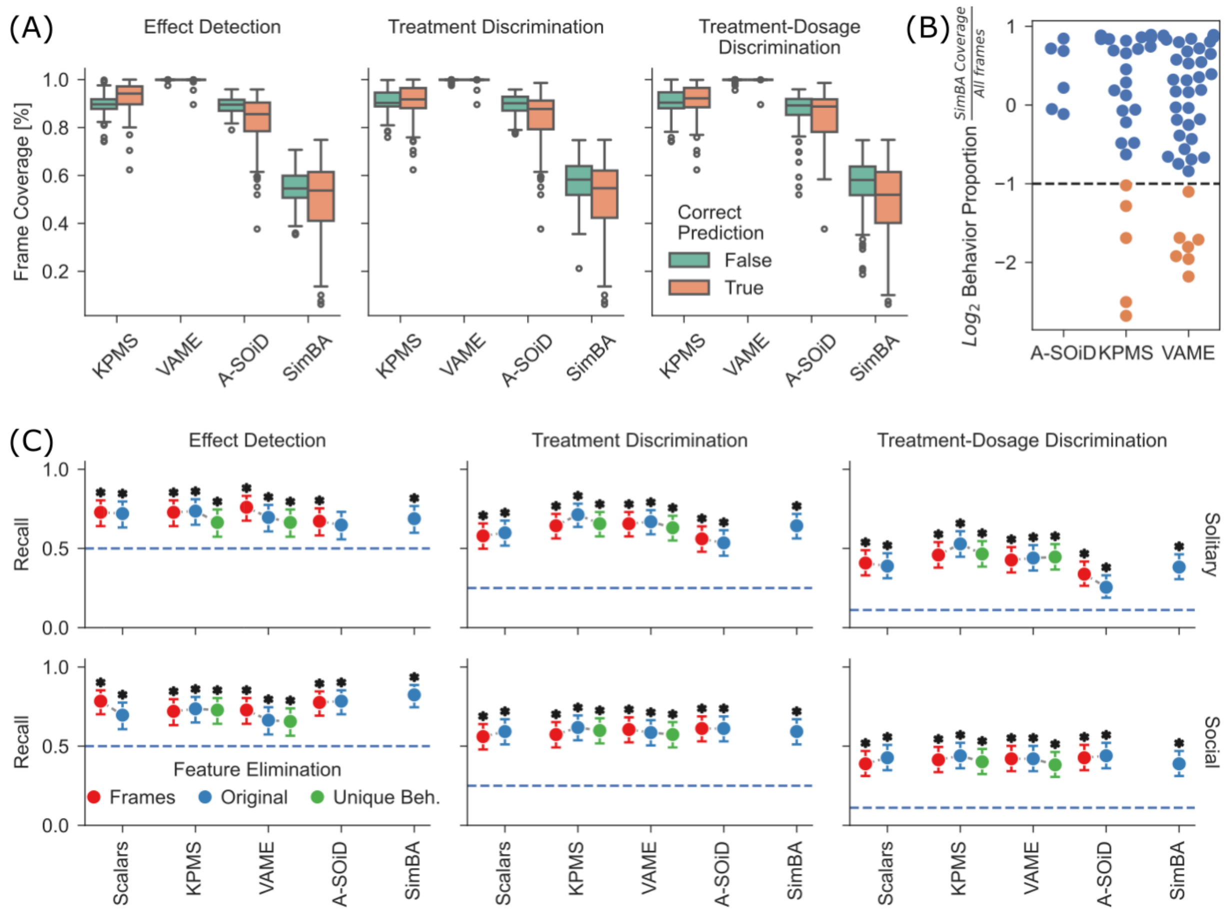


**Suppl. Figure 4. Overview over differences in recording coverage and unsupervised segmentation specific behaviors.** (A) Comparison of label coverages across models and analytical levels. Coverage is compared between correct and incorrect classifications. No statistical tests were run. (B) Swarmplot of log2 proportion ratios for each behavior within 3 behavioral summary models, calculated between recording frames covered by SimBA and those not labeled by SimBA. Behaviors with a twofold reduced frequency within the SimBA labeled frames were marked and will be referred to as “unique behaviors” of the unsupervised segmenters. (C) Global recall across tasks, context, and behavioral summary models. Input data to the classification was modified by dropping frames that were not covered by SimBA’s labels in all other models, and by dropping behaviors detected to be unique to the unsupervised models. The outcome is compared against the unmodified dataset. Same tests and annotation as in Figures 4-6.

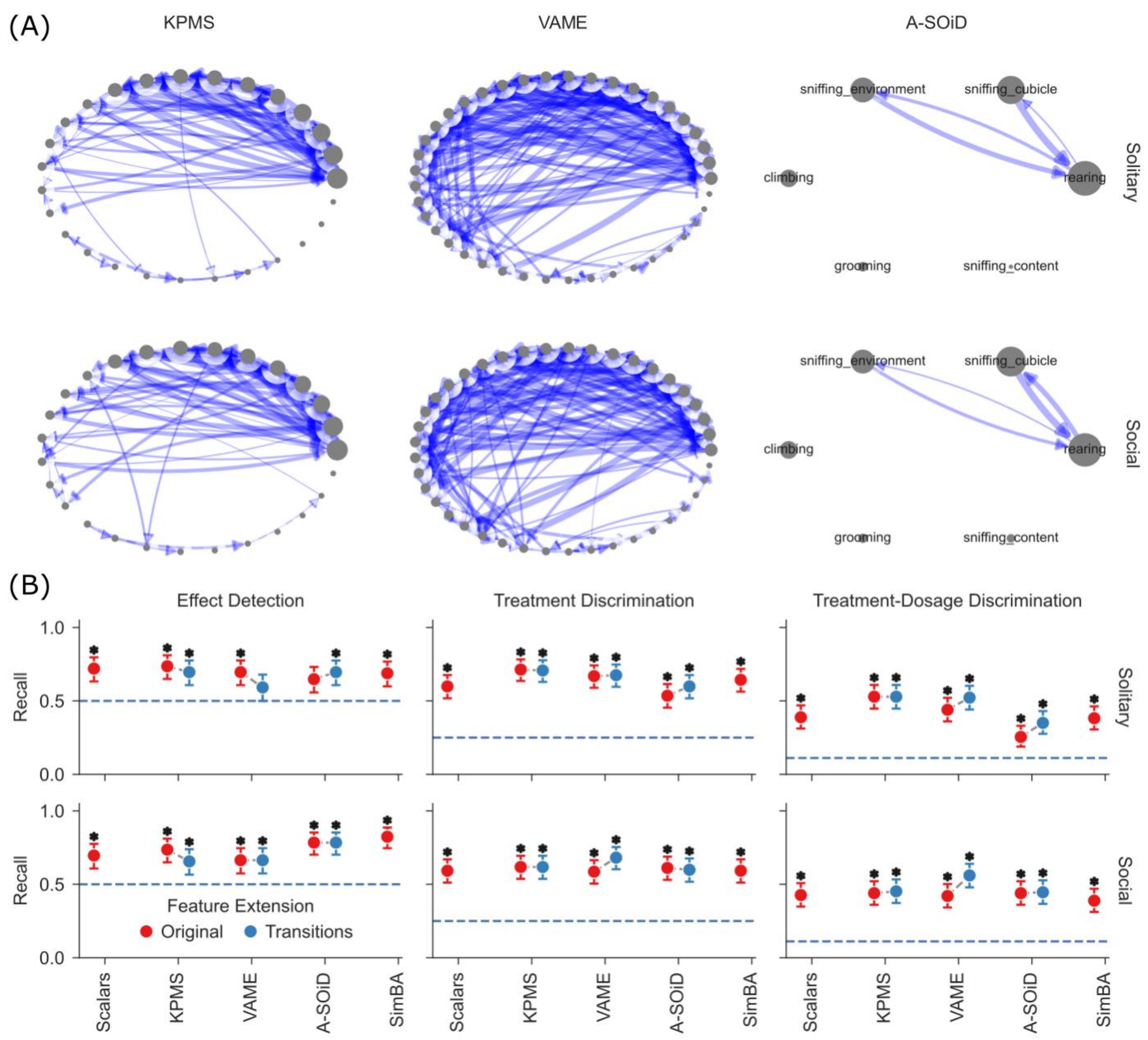


**Suppl. Figure 5. Overview over transitions between continuous model labels.** (A) Circular transition networks visualizing the qualitative transition probabilities of the top 10% most frequent transitions across continuous models and contexts. Circle size indicates frequency of behavioral labels, edge width the frequency of transitions. Named labels were only included for the supervised A-Soid model. (B) Global recall across tasks, contexts, and behavioral summary models. Input data to the classification was modified by adding transition proportions to the feature vector of the continuous models. The outcome is compared against the unmodified dataset. Same tests and annotation as in Figures 4-6.

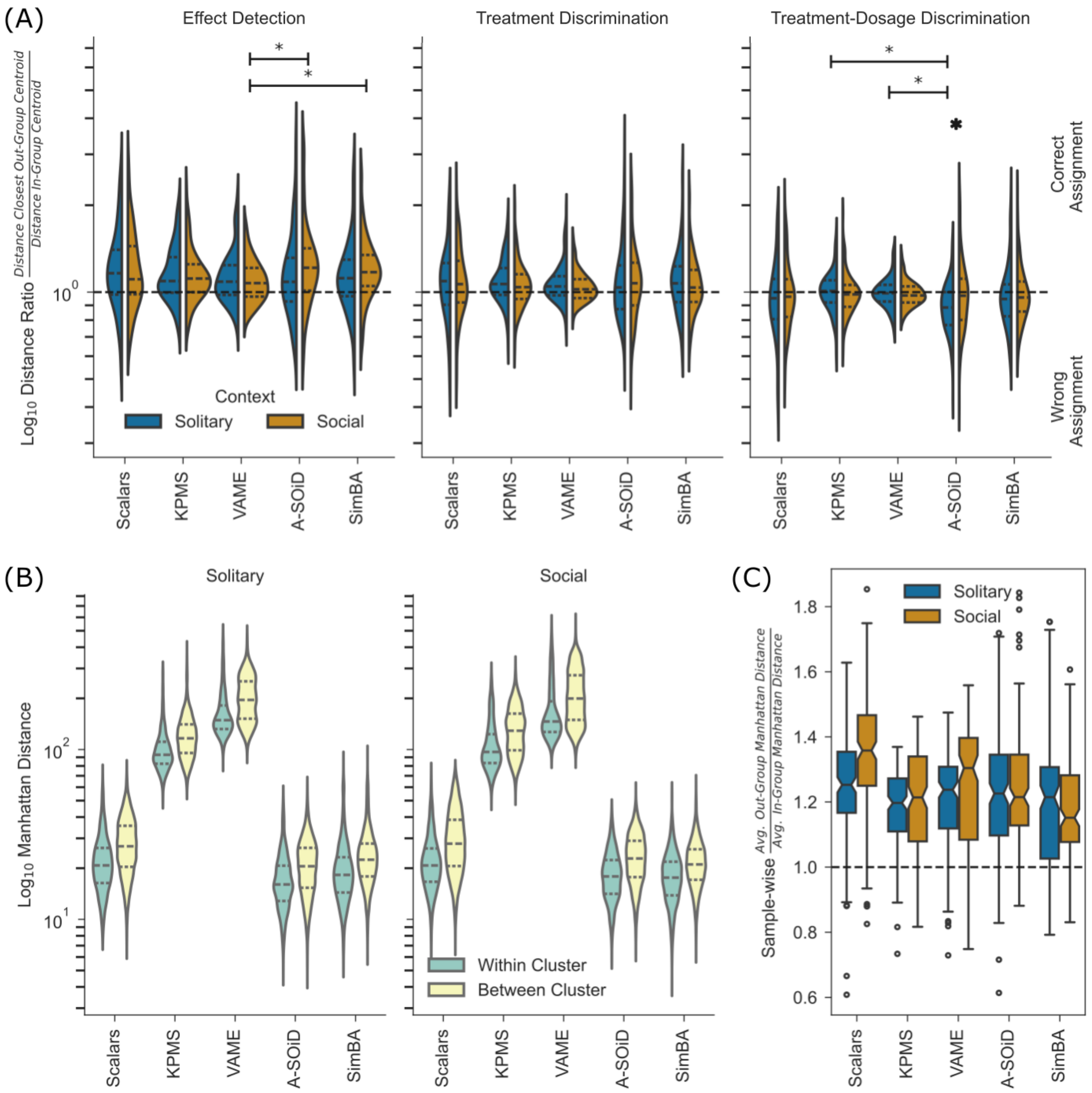
(A)
Effect Detection
Treatment Discrimination
Treatment-Dosage Discrimination
Log10 Distance Ratio Distance Closest Out-Group Centroid / Distance In-Group Centroid
10^0
Context
Solitary
Social
Correct Assignment
Wrong Assignment
Scalars
KPMS
VAME
A-SOiD
SimBA
(B)
Solitary
Social
Log10 Manhattan Distance
10^2
10^1
Within Cluster
Between Cluster
(C)
Sample-wise Avg. Out-Group Manhattan Distance / Avg. In-Group Manhattan Distance
1.8
1.6
1.4
1.2
1.0
0.8
0.6

**Suppl. Figure 6. Alternative distance analysis and comparison of within- and between group variances across models.** (A) Violinplot of distance ratios for each recording. Distance ratios were calculated based on the quotient of the distance to the closest centroid belonging to a different group, over the distance to the centroid belonging to the group of the sample. Ratios above 1 indicate a correct prediction, as the samples true group centroid is closer than the next closest centroid. Comparisons were run within models and across context, and within context and across models. Significant differences are indicated by an asterisk or bar and asterisk (Mann Whitney U Test, Bonferroni correction, p<= 0.05). (B) Violinplots of Manhattan distances within groups, and between groups. Distances are calculated pairwise between all samples and plotted separated by context and model. Statistical tests were run comparing within and between group distances, and per model and context, all were significant (Mann Whitney U Test, Bonferroni correction, p<=0.05). (C) Boxplots of ratio of average between-group distance over average within-group distance, calculated per sample and plotted across models and contexts. A line at ratio=1 (same average distance within and between groups) was added to facilitate comparisons. No statistical tests were run.

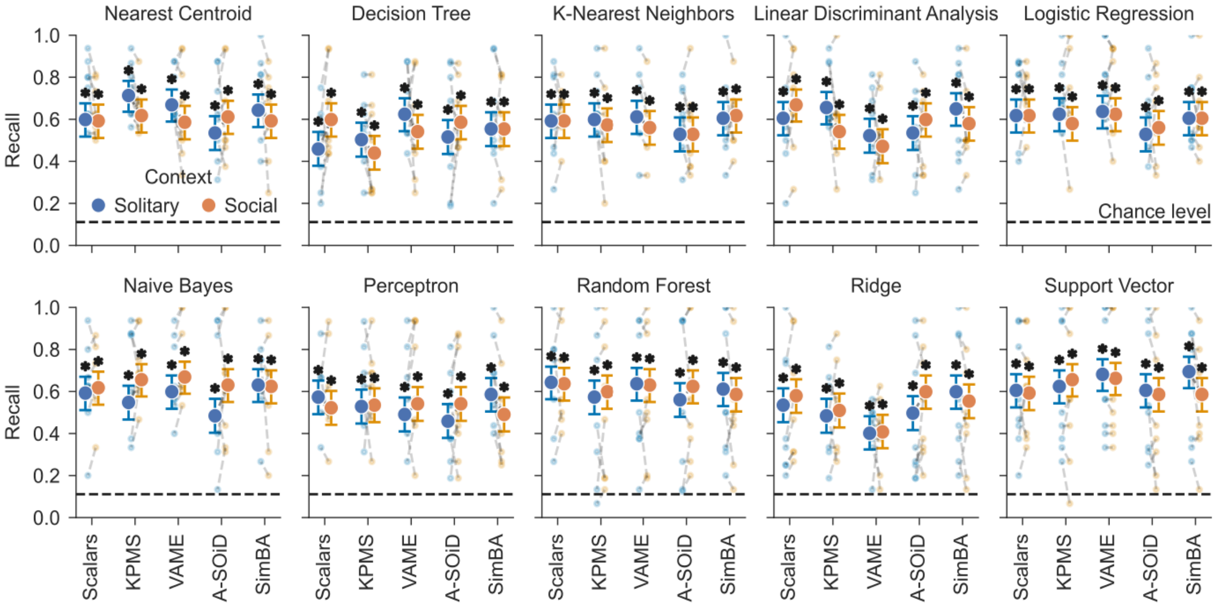


**Suppl. Figure 7. Outcome of various alternative classifier models.** 10 classifiers were applied to the features returned by the behavioral summary models. Same tests and annotation as in Figures 4-6.

14

**Suppl. Table 1.** ***Table of single features leading to discrimination performance above chance level***. Every feature shown corresponds to a single pre-processed output from a behavioral summary model (rows) leading to performance significantly above chance in discriminating at a certain level (sub columns) during a particular phase (top columns). For details on the discrimination tasks see Results. Numbers in brackets correspond to global recall (across all samples) in the given task.

| **Context** | **Solitary** | | | **Social** | | |
|---|---|---|---|---|---|---|
| **Model \ Task** | vehicle | drug-only | drug-dose | vehicle | drug-only | drug-dose |
| **Locomotion** | | moved_distance_m_None (0.54), body_length_std_mm_Mouse (0.5), body_length_std_mm_Object (0.48) | moved_distance_m_None (0.3) | | moved_distance_m_None (0.46) | |
| **KPMS** | right_turn_and_sniff_None (0.91), left_turn_and_forward_Mouse (0.78), cubicle_rear_None (0.77) | climb_stretch_on_right_side_Mouse (0.5), climb_stretch_on_left_side_Mouse (0.49), cubicle_rear_Object (0.49), unclear_head_right_None (0.48), unclear_head_right_Object (0.47), cubicle_rear_Mouse (0.46) | left_turn_and_forward_Mouse (0.29), left_turn_Mouse (0.28) | right_turn_and_sniff_None (0.89), cubicle_rear_None (0.82), climb_start_None (0.78), climb_head_left_Mouse (0.76) | climb_stretch_on_right_side_Mouse (0.5), climb_stretch_on_left_side_Mouse (0.47), right_turn_Mouse (0.47) | targeted_sniff_None (0.28) |
| **VAME** | | rearing_Mouse (0.55), climb_or_groom_right_turn_Mouse (0.5), climbing_turn_left_Object (0.5), head_turn_right_None (0.49), rearing_Object (0.49), climbing_face_upwards_Mouse (0.49), sniff_with_right_turn_Mouse (0.48), move_forward4_Object (0.48), move_and_stop_None (0.48), unclear_forward_None (0.48) | move_and_stop_None (0.31), sniff_with_left_turn_Mouse (0.3) | move_forward4_None (0.8), move_and_stop_None (0.8), climbing_face_upwards_None (0.78) | rearing_Object (0.48), unclear_forward_None (0.46), climbing_turn_left_Mouse (0.46) | rearing2_None (0.32), move_and_stop_None (0.3) |
| **A-SOiD** | | sniffing_cubicle_Mouse (0.48), climbing_Mouse (0.46) | | sniffing_content_Mouse (0.82), grooming_Object (0.78), sniffing_content_Object (0.77) | sniffing_content_Object (0.46), climbing_Object (0.46) | sniffing_cubicle_Mouse (0.31) |
| **SimBA** | grooming_Object (0.79) | rearing_None (0.5), climbing_Mouse (0.47) | | | climbing_Object (0.49) | rearing_None (0.29), sniffing_cubicle_Object (0.28) |
