## Supplementary Methods and Materials for "Comparison of multiple video tracking-based behavioral summary approaches for compound discrimination"

#### 1 Video recording gaps

Analysis of the video recordings revealed short breaks of various lengths (1 to 47 frames per occurrence, 1 to 773 cumulative frames in 1 to 42 separate occurrences per video) in a small subset of videos (46 out of 320, ~14.4%). As each video was 10 minutes long, this affected only a small part of the total dataset (10921 frames / 7.21 minutes out of roughly 4844160 frames / 53.3 hours, ~0.23%). Relatively more frames were affected in the Vehicle and seltorexant datasets (~0.34% and 0.42% respectively) compared to amphetamine and modafinil recordings (~0.09% and ~0.16% respectively). We resolved this issue by removing affected videos entirely from the training dataset and only applying trained models to the full dataset (including affected videos). This resulted in our training dataset of only unaffected Vehicle treatment recordings containing 49 videos (~8.2 hours of recording, out of a potential 64 videos, 10.7 hours). All videos were used for inference and downstream analysis.

#### 2 Further Caveats

It is important to note that the choice of hyperparameters is essential to the performance of all models shown in this study. For VAME and KPMS, we maintained the same workflow already established in our earlier study with KPMS (Ritter et al. 2025) and are thus confident that the selected parameters lead to reproducible and successful extraction of motifs. The selection of hyperparameters was guided by the existing documentation of both tools (see Supplementary Methods), and the application was done in notebooks packaged with the selected releases.

While a reduced tracking dataset was used to train the A-SOiD and SimBA models, their recall does not differ significantly from the unsupervised methods. A-SOiD's training data was further halved to allow for the application of active learning on the same data as the other tools. While it is possible to correct both caveats, this would not change the primary result of the study.

Our choice of parameters to extract in the parametric aggregation model was guided by an experienced subject matter expert in our lab, inspired by parameters extracted with Stoelting Any-Maze (Figueiredo Cerqueira et al. 2023), and is thus subjective and possibly hard to generalize to other labs. We acknowledge this limitation and tried to select measures that could be easily generated by established tracking tools commonly used in other groups (e.g. Stoelting ANY-Maze).

#### 3 Keypoint formats

| Index | Unsupervised Schema | Dropped? | SimBA Schema |
|---|---|---|---|
| **1** | nose | | Nose |
| **2** | head | **x** | |
| **3** | ear_left | | Ear_left |
| **4** | ear_right | | Ear_right |
| **5** | spine_cervical | **x** | |
| **6** | spine_thoratic | | Center |
| **7** | side_left | | Lat_left |

| | | | |
|---|---|---|---|
| **8** | side_right | | Lat_right |
| **9** | spine_lumbar | **x** | |
| **10** | tail_base | | Tail_base |
| **11** | tail_center | **x** | |
| **12** | tail_tip | | Tail_end |

#### 4 Model configurations

As most parameters were unchanged from the default settings (see Methods for versions), we will only list those that we adapted manually for our dataset.

##### 4.1 Keypoint-MoSeq

| **Category** | **Parameter** | **Value** | **Comments** |
|---|---|---|---|
| ***FITTING*** | conf_threshold | 0.3944923929354759 | noise_calibration widget |
| ***HYPER PARAMS*** | intercept | 0.1682062564155024 | “ |
| | slope | -1.071950416420913 | “ |
| | latent_dim | 4 | Determined through PCA |

##### 4.2 VAME

| **Category** | **Parameter** | **Value** | **Comments** |
|---|---|---|---|
| **Preprocessing** | pose_confidence | 0.3944923929354759 | value from KPMS |
| ***RNN model general hyperparameter*** | time_window | 13 | Reduced for 25.23 Hz video |
| | prediction_steps | 7 | “ |
| ***Segmentation*** | n_clusters | 100<br>41 | … for test<br>… for inference |

##### 4.3 BORIS

| **Behavior / excludes** | **Climbing** | **Grooming** | **Rearing** | **Sniffing (Cubicle)** | **Sniffing (Content)** | **Sniffing (Environment)** |
|---|---|---|---|---|---|---|
| **Climbing** | | **x** | **x** | | | **x** |
| **Grooming** | **x** | | **x** | **x** | **x** | **x** |
| **Rearing** | **x** | **x** | | | | |
| **Sniffing (Cubicle)** | | **x** | | | | **x** |
| **Sniffing (Content)** | | **x** | | | | **x** |
| **Sniffing (Environment)** | **x** | **x** | | **x** | **x** | |

#### 4.4 SimBA

| Category | Parameter | Value | Comments |
|---|---|---|---|
| **SML settings** | **no_targets** | 6 | |
| | **target_name_1** | grooming | |
| | **target_name_2** | climbing | |
| | **target_name_3** | rearing | |
| | **target_name_4** | sniffing_content | |
| | **target_name_5** | sniffing_cubicle | |
| | **target_name_6** | sniffing_environment | |
| ***threshold_settings*** | **threshold_1** | 0.354 | Calculated for maximum F1 |
| | **threshold_2** | 0.538 | “ |
| | **threshold_3** | 0.4225 | “ |
| | **threshold_4** | 0.2795 | “ |
| | **threshold_5** | 0.3955 | “ |
| | **threshold_6** | 0.318 | “ |
| ***Minimum_bout_lengths*** | **min_bout_1** | 277 | Calculated from bottom 5$^{th}$ percentile of training data |
| | **min_bout_2** | 2339 | “ |
| | **min_bout_3** | 476 | “ |
| | **min_bout_4** | 198 | “ |
| | **min_bout_5** | 199 | “ |
| | **min_bout_6** | 199 | “ |

#### 4.5 A-SOiD

| Category | Parameter | Value | Comments |
|---|---|---|---|
| **Project** | **MULTI_ANIMAL** | True | Required for bottom-up SLEAP model |
| **Processing** | **LLH_VALUE** | 0.3944923929354759 | value from KPMS |
| | **MIN_DURATION** | 0.08 | Corresponding to 3 frames, based on minimum length found in training labels |
| | **MAX_SAMPLES_ITER** | 70 | Based on training dataset size |
| | **N_SHUFFLED_SPLIT** | None | |

### 5 Unsupervised syllable names

#### 5.1 Keypoint-MoSeq syllables

| Syllable | Label | Bout Frequency [%] | Short description |
|---|---|---|---|
| **0** | targeted_sniff | 11.97 | 20/24 show some form of sniffing in a corner or at/behind a cubicle, maybe more related to head stretch? Most likely strongly related to immobility |
| **1** | body_raise | 11.98 | 21/24 show a component of rearing, raising the body. Can also apply to stopping forward movement (9/24), potentially related to 2D recording. |
| **2** | right_turn | 8.64 | 22/24 show rightward turn in different situations (rearing, movement, sitting). Can also occur as part of rearing sniffing. |
| **3** | left_turn | 8.86 | 20/24 show leftward turn, occasionally from rearing (9/24) |
| **4** | unclear_contraction_head_down | 6.31 | 13/24 quite unclear head downwards movement maybe combined with body contraction, maybe related to rightwards turn (15/24) |
| **5** | short_dart | 6.56 | 17/24 related to (short) movement forward, often out of stationary behavior (15/24) |
| **6** | head_up | 5.19 | 21/24 related to upwards movement of head, often related to (beginning) rearing (17/24) |
| **7** | unclear_head_retract_left | 5.55 | 15/24, short leftwards turn of head, not as clearly defined as previous syllables |
| **8** | right_turn_and_forward | 6.08 | 21/24, mouse turns right and moves forward |
| **9** | left_turn_and_forward | 6.03 | 21/24, mouse turns left and moves forward |
| **10** | unclear_head_right | 1.98 | 14/24, a very short head turn to the right |
| **11** | head_left | 2.07 | 20/24, leftwards version of 10, sometimes combined with short movement forwards (11/24), like 9 |
| **12** | assisted_body_raise | 2.0 | 19/24, move into rearing along cubicle or walls |

| | | | |
|---|---|---|---|
| **13** | rightwards_rear | 1.58 | 19/24, related to rearing with rightwards head/body turn |
| **14** | dart_stop | 1.54 | 17/24 related to second half of forward movement, slowing down |
| **15** | cubicle_rear | 1.19 | 20/24 show rightwards oriented rearing or climbing on cubicle, 20/24 in contact with cubicle |
| **16** | climb_left | 0.97 | 21/24 show left-oriented climbing behavior, 24/24 are climbing on cubicle |
| **17** | climb_start | 0.96 | 16/24 begin to climb on cubicle, other show rearing, or complicated climbing behavior or drops |
| **18** | right_turn_and_sniff | 0.82 | 15/24 show a rightward turn of some kind followed by a sniffing motion, usually close to the cubicle (17/24) |
| **19** | climb_right | 0.74 | 21/24 show rightwards turn during climbing, 24/24 are climbing on cubicle |
| **20** | climb_stretch_on_right_side | 0.65 | 21/24 show some climbing related behavior, looks like stretches are frequent, but always on right side of cubicle |
| **21** | climb_stretch_on_left_side | 0.53 | 17/24 show similar, diverse climb-related behavior as 20, but this time on left side of cubicle |
| **22** | climb_head_left | 0.53 | 20/24 turn head leftwards while climbing, 17/24 are on top or on the top half of the cubicle |
| **24** | climb_left_sniff | 0.6 | 22/24 turn left while climbing, most (20/24) seem to sniff cubicle while doing so |
| **25** | cubicle_corner_sniff_right | 0.62 | 17/24 sniff the right corner between cubicle and wall |
| **26** | climb_down_left | 0.51 | 21/24 climb down from the cubicle, while turning/turned left |
| **30** | unclear_dart_possibly_stop | 0.5 | 14/24 show some forward movement, quite unclear, might be related to stopping |

##### 5.2 VAME

| Motif | Label | Bout Frequency [%] | Short description |
|---|---|---|---|
| **0** | move_forward | 3.47 | Moving forward, sometimes into rearing (14/24) |
| **1** | stretch_or_rear | 4.34 | Rearing or stretching associated (15/24) |
| **2** | climb_or_groom_shorten | 1.81 | Related to climbing and grooming, potentially due to shorteing of body relative to camera (16/24) |
| **3** | head_turn_right | 4.58 | various right head turns, some from rearing, some on ground (11/24) |
| **4** | turn_left_and_move | 3.56 | Movement forward with leftward turn (16/24) |
| **5** | move_and_stop | 1.13 | Usually short movement followed by shortening of body and stopping, so either sniffing at walls or general sniffing(13/24) |
| **6** | sniff_with_left_turn | 3.7 | Sniffing at cubicles and walls and left turn (20/24) |
| **7** | turn_right_stationary | 0.55 | Stationary turn rightwards, sometimes part of grooming (19/24) |
| **8** | turn_right | 3.99 | various movements towards the right, with varying contexts, sometimes grooming (13/24) |
| **9** | turn_left_and_move2 | 1.75 | Left turn followed usually by movement (13/24) |
| **10** | sniff_with_left_turn2 | 2.72 | Sniffing at cubicles and walls and left turn (17/24) |
| **11** | turn_left | 2.16 | Leftwards turn, sometimes with forwards movement (13/24) |
| **12** | move_forward_and_up | 3.28 | Forward movement sometimes leading into an upwards movement into rearing (16/24) |
| **13** | climb_or_groom_right_turn | 0.55 | Turning right during climbing or grooming (17/24) |
| **14** | sniff_with_right_turn | 0.69 | Sniffing with general rightwards orientation, or actual turning (18/24) |
| **15** | move_forward2 | 3.08 | General forward movement (15/24) |
| **16** | sniff_or_climb | 1.85 | Sniffing on floor or during climbing, on ground leftwards bias, during climbing rightwards (19/24) |
| **17** | head_raise | 3.22 | General movement of head upwards, often following forward movement (14/24) |

| | | | |
|---|---|---|---|
| **18** | turn_left2 | 2.08 | Leftwards turn, often related to rearing (17/24) |
| **19** | turn_left3 | 2.87 | General turn leftwards (18/24) |
| **20** | turn_right2 | 1.42 | Right turn, sometimes followed by moving forward, sometimes during grooming (14/24) |
| **21** | move_forward3 | 3.36 | General forward movement (16/24) |
| **22** | rearing | 1.04 | General movement related to rearing (15/24) |
| **23** | up_into_rear | 3.0 | Raising head, with upwards movement into rearing (15/24) |
| **24** | sniff_with_right_turn2 | 2.92 | Sniffing with head down, with rightwards turn (18/24) |
| **25** | turn_right_and_move | 1.81 | Right turn followed often by forward or upward movement (16/24) |
| **26** | head_down_and_sniff | 2.9 | Moving head slightly down towards sniffing, often with rightwards bias (14/24) |
| **27** | turn_right_and_move2 | 3.36 | right turn followed often by movement forward (16/24) |
| **28** | head_right | 0.73 | Turning head right (18/24) |
| **29** | rearing2 | 3.34 | Rearing-related, with shortening? Shows both rearing start and end (17/24) |
| **30** | rearing3 | 2.0 | Rearing-related, again both start and end shown (14/24) |
| **31** | move_forward4 | 0.72 | general movement forward (15/24) |
| **32** | head_turn_left | 2.75 | Head turn leftwards (19/24) |
| **33** | climbing | 0.93 | General climbing related motif, generally not much turning (23/24) |
| **34** | sniffing_ground | 2.99 | Sniffing on floor generally (14/24) |
| **35** | climbing_face_upwards | 1.16 | Climbing while facing upwards, some rearing (20/24) |
| **36** | head_up_from_down | 4.47 | Raising head form ground, usually rightwards bias (17/24) |
| **37** | climbing_turn_left | 0.72 | During climbing or sometimes rearing, body slightly bent left (15/24) |
| **38** | sniffing_ground2 | 3.48 | Sniffing on ground, usually at corners, such as walls or cubicles (14/24) |
| **39** | head_raise_right | 2.43 | Raising head, with slight rightwards bias (19/24) |
| **40** | unclear_forward | 2.98 | Unclear motif showing often forwards movement, with occasional result in sniffing or stopping (13/24) |
